# The developing leaf of the wild grass *Brachypodium distachyon* at single-cell resolution

**DOI:** 10.1101/2025.11.03.686118

**Authors:** Lea S. Berg, Paola Ruiz Duarte, Inés Hidalgo Prados, Nathan T. Lacombe, Rashmi Tandon, Isaia Vardanega, Jan E. Maika, Roxane P. Spiegelhalder, Ambuj Gore, Heike Lindner, Rüdiger Simon, Michael T. Raissig

**Author notes:** Corresponding author: MTR.

## Abstract

Leaves are the plant’s main photosynthetic organs and drive Earth’s primary production. Grasses form longitudinal leaves with parallel venation and graminoid stomata. Yet, how distinct leaf tissues coordinatively develop to build functional grass leaf anatomy is not well understood. Here, we decoded the developing grass leaf from vegetative meristems to mature tissues using single-cell RNA-sequencing in the wild grass *Brachypodium distachyon*. In-depth analysis of epidermal clusters and multiplexed whole-mount RNA-fluorescence *in situ* hybridization resolved most epidermal lineages and confirmed them *in planta*. Gene regulatory network analysis distinguished the targetome of the two co-expressed, yet functionally divergent stomatal transcription factors *BdMUTE* and *BdFAMA*. Finally, we used our dataset to identify and functionally describe *BdGRAS32*’s role in cell division inhibition and a stomata-specific function for a cell wall modifying enzyme. Together, our single-cell grass leaf atlas enables the dissection of developmental processes that shape the leaf sustaining global food production.

## Introduction

More than 40% of the Earth’s surface is dominated by grasses^1^ and the three cereal crops maize, rice, and wheat directly produce more than half of the calories consumed by humans^2^. Thus, the grass leaf is the photosynthetic powerhouse that fuels ecosystems and our civilization alike. The longitudinal sheathing leaf of grasses and other monocots is an evolutionary innovation, where the leaf sheath protects the shoot apical meristem and developing leaves^3^. The parallel venation provides additive water transport capacity^4^ and morphological improvements to the graminoid stomata^5,6^ enable fast and water-use-efficient stomatal aperture control^7–9^.

Yet, despite the innovative nature and humanitarian importance of the grass leaf, little is known regarding the processes that generate and coordinate the distinct leaf tissues and cell types to build a functional photosynthetic organ. This is partly due to difficulties inherent to grasses as model systems like their size, their massive genomes and their resistance to being transformed by *Agrobacterium tumefaciens*^10^. As a consequence, a lack of tissue-specific reporter lines prevented the description of the transcriptomic signatures of distinct grass leaf tissues and cell types using cell sorting and bulk RNA-sequencing approaches as was done in the eudicot model system *Arabidopsis thaliana* (e.g. ^11–13^). In recent years, the advance of single-cell RNA-sequencing (scRNA-seq) facilitated the generation of transcriptomes at single-cell resolution and clustering of cells with similar transcriptomic signatures^14^. These new technologies do not require the isolation of specific cell types or tissues using laser microdissection or fluorescent-activated cell sorting (FACS) followed by bulk RNA-seq. Instead, dissociation of tissues and pairing of single plant cells (protoplasts) with beads containing bead-specific barcodes (either in droplets or plates) allows for the generation of single-cell cDNA libraries, massive parallel sequencing of single cell populations and rapid demultiplexing of reads into single-cell transcriptomes. Strikingly, single-cell transcriptomics in plants repeatedly showed the capability to detect most if not all expected cell types of a given organ and, in addition, revealed cellular heterogeneity and developmental transitions within a tissue of diverse plant species (e.g.^15–33^).

Despite the generation of single-cell atlases in (domesticated) grasses, their resistance to transformation remains a significant problem to confirm the clusters identified *in silico* also *in planta* with reporter genes or to generate CRISPR/Cas9-mediated gene knock-outs of cluster-specific genes with a putative regulatory function. We therefore work with the wild Pooideae model grass *Brachypodium distachyon* (Fig. 1A). Not only is *B. distachyon* small in size and easy to grow with limited space, it is also quite amenable to *A. tumefaciens*-mediated transformation compared to most other grasses, even without access to large-scale transformation facilities. Furthermore, research on a wild grass likely reveals higher gene diversity, as wild grasses did not suffer from domestication-induced loss of genetic diversity^34^.

**Figure 1.**
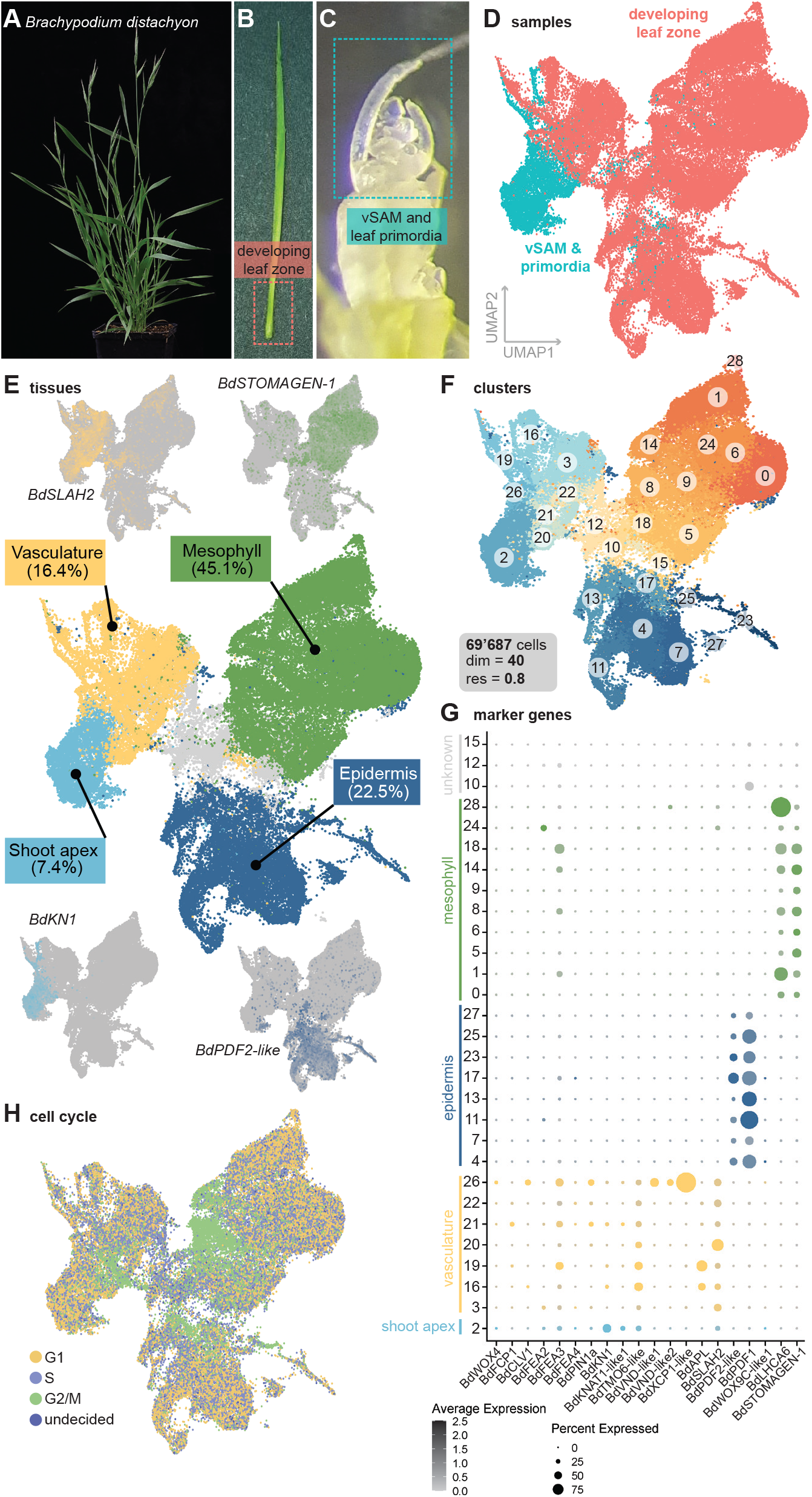
Single-cell atlas of the developing *B. distachyon* leaf from vegetative shoot apex to mature tissues. **(A)** *Brachypodium distachyon* flowering plant. **(B)** Young, pulled-out, and not yet unrolled leaf used to isolate the leaf developmental zone (red box). **(C)** Dissected shoot apex with vegetative shoot apical meristem (vSAM), leaf primordia and young outgrowing leaves. Collected tissue is highlighted with a blue box. **(D)** Whole dataset UMAP plot indicating the tissue-origin of the cells (vSAM and primordia in blue, developing leaf zones in red). **(E)** Whole dataset UMAP plot with tissue annotations (percentage of total cells in brackets) and exemplary marker gene feature plots next to each tissue annotation. N = 69’687 cells. **(F)** Whole dataset UMAP plot with numbered Seurat clusters. Number of total cells, dimensions and resolution are indicated. **(G)** Dot plot showing expression of marker genes in the clusters. Dot size represents the percentage of cells within a cluster that express the gene and color saturation represents expression strength. Tissues are color-coded as in (E). **(H)** Whole dataset UMAP plot with color indicating the cell cycle stage of each cell. Also see Fig. S1 and S2.

Grass leaf development is initiated at the vegetative shoot apical meristem (vSAM). In the emerging primordium, the stem cell factor *KNOTTED1* (*KN1*) is downregulated, auxin maxima contribute to primordium outgrowth^35–38^, and a WUSCHEL-HOMEOBOX module drives medio-lateral leaf expansion^29,39^. In leaf primordia, the ground meristem forms the vasculature and mesophyll and the protodermal stem cells form all epidermal cell types^40^. In the epidermis, protodermal stem cells initially proliferate symmetrically before an asymmetric, transverse patterning division forms a small apical and a larger basal cell^41–43^. Mediolateral patterning of the semi-clonal epidermal cell files is determined by antagonistic transcription factor (TF) programs. Grass SPEECHLESS (SPCH) homologs together with INDUCER OF CBF EXPRESSION1 (ICE1) determine stomatal cell file identity^44–46^, whereas SQUAMOSA PROMOTER BINDING PROTEIN-likes (SPLs) establish hair cell (HC) file identity^47,48^. In HC files, the apical HC precursors directly form prickle hair cells or macrohairs^49^. In stomatal files, the apical guard mother cells (GMCs) elongate and laterally deploy the mobile grass MUTE homolog to establish subsidiary mother cell (SMC) identity in neighboring cells^9,50^. SMCs then polarize and an asymmetric longitudinal division forms the proximal subsidiary cell (SC)^51,52^. Finally, a symmetric longitudinal division that involves cell-autonomous MUTE function forms the guard cell (GC) pair^53^, before FAMA and SCREAM 2 (SCRM2) induce the stomatal pore and guide dumbbell GC morphogenesis^44,45,54,55^.

Importantly, even after primordium outgrowth, developing grass leaves maintain a spatiotemporally separated developmental leaf zone, where all divisions, cell elongations and morphogenetic processes occur in a strict base-to-tip developmental gradient. Here, we leveraged this peculiarity of grass leaf development to isolate developing leaf zones of 3 weeks old *B. distachyon* plants, which were dissociated and single cell transcriptome libraries were built using the droplet-based 10X Genomics platform. In addition, we isolated shoot apices including vSAMs and early leaf primordia. Together, we generated a single-cell atlas from shoot apex to mature grass leaf cell types consisting of almost 70k cells. Marker gene analysis readily identified the four major tissues (i.e., shoot apical stem cells, vasculature, mesophyll and epidermis). *In silico* isolation and re-analysis of the epidermal cluster identified all cell lineages of the abaxial epidermis. Multiplexed whole-mount RNA fluorescence *in situ* hybridization (FISH) confirmed the identified clusters *in planta*. Subsetting of the stomatal lineage and pseudotime analysis of the developing GCs solidly recapitulated previously described developmental modules and identified novel, developmental-stage-specific marker genes. Analysis of homologs of previously reported genetic programs involved in epidermal patterning allowed to pinpoint clusters that will produce pavement cells, a cell type that is widely unexplored. Single-cell gene regulatory network (GRN) analysis of the stomatal lineages revealed specific targetomes of the co-expressed yet functionally divergent TFs BdMUTE and BdFAMA in GMCs indicating that our atlas can resolve cell-type- and developmental stage-specific GRNs. As a proof-of-concept, we identified a role for *BdGRAS32* in dividing leaf cells and suggested a role for *BdPME53-like*-mediated processes in stomatal patterning. Together, our high-quality dataset will facilitate the identification of marker genes and GRNs in the developing leaf of *B. distachyon* and can serve as a comparative single-cell gene expression resource for research on both wild and domesticated grass leaves.

## Results

### A single-cell transcriptomic atlas of the developing grass leaf

While some genes involved in the development of the grass leaf have already been investigated^40^, much remains unexplored regarding the genetic programs that form and coordinate the diverse cell types of the leaf. Therefore, we generated a single-cell transcriptomic atlas of shoot apices and developing leaves using the wild grass model *Brachypodium distachyon* (Fig. 1A) to identify genetic modules that accommodate the development of distinct grass leaf tissues and cell types.

We harvested two different tissue samples. First, young, not yet unrolled leaves were extracted from the embracing sheath of the older leaf and the lowest 3-5 mm were collected for seven independent “leaf developmental zone” libraries. Owing to the strict acropetal (base-to-tip) developmental gradient and clearly separated developmental leaf zone, these samples likely contained all developmental stages and tissues of the leaf (Fig. 1B, Table S1). In addition, we dissected vegetative shoot apices including young leaf primordia and generated two independent “vSAM and primordia” libraries (Fig. 1C, Fig. S1C, Table S1). Then, cell wall digestion released protoplasts (Fig. S1A,B), which were then used for droplet-based 10X Genomics single-cell RNA-sequencing (scRNA-seq). The resulting datasets, generated at different time points and locations (Table S1), were demultiplexed using Cell Ranger and subsequently filtered to exclude cells with high ambient RNA content, high mitochondrial (>5%) or chloroplast (>10%) RNA content, too few or too many genes and transcripts detected (<1250 and >50’000 unique molecular identifiers (UMIs); <500 and >10’000 features) and removed doublets to generate a high-quality dataset (see STAR Methods). All nine datasets were then merged to create the final dataset containing 69’687 cells (expressing 35’584 out of 39’068 genes, Table S1, Table S2; see STAR Methods). Little batch effects were observed (Fig. S1C) and, thus, batch effect correction through integration was deemed unnecessary as it would have removed the biologically inherent differences between shoot apex and leaf developmental zone. Even without integration, there was considerable overlap between these two sample types, nicely re-creating the developmental trajectories from shoot apex to mature leaf tissues (Fig. 1D).

After dimensionality reduction and uniform manifold approximation and projection (UMAP) clustering (Fig. 1F), expression patterns of marker gene homologs were used to assign the clusters to shoot apices, vasculature, mesophyll and epidermis (Fig. 1E-G, Fig. S1D-W, Table S3). There was considerable overlap between the vasculature and the shoot apex possibly because the vasculature develops first and originates from the inner ground meristem that constitutes most of the shoot apex and leaf primordia^40^ and/or because more mature vascular cells did not protoplast well or were filtered out due to cell size limits of our protocol (Fig. 1E,G, Fig. S1D-R). We were, however, able to detect distinct expression patterns of shoot apex and early vasculature markers (Fig. 1E,G, Fig. S1D-R). A recently published reporter of the barley ortholog of *Bd-FON2-LIKE CLE PROTEIN 1* (*BdFCP1*)^56^ that showed expression in the vSAM and leaf primordial epidermis helped to assign tissue identities (Fig. 1E,G, Fig. S1E, Fig. S2E,G). Additionally, spatial transcriptomics and imputed data of stem cell homologs in barley further resolved stem cell clusters (Fig. S2A-F,H-K; ^27^). *HvCLV1* marks outer vSAM layers (^56^; Fig. S1F, Fig. S2A,B, also seen in inflorescence meristems in the maize ortholog *ZmTD1*^57^) and *CRABS CLAW* (*BdCRC*, in barley *HvCRC*) helped to indicate leaf primordial tissues (^56,58^; Fig. S2C,D). *PIN-FORMED 1a* (*BdPIN1a, HvPINa*) and *PIN1b* (*BdPIN1b, HvPIN1/HvPIN1b*) pinpointed leaf primordia and inner meristem, respectively (Fig. S2H-K; ^59,60^). The expression pattern of a *B. distachyon* transcriptional reporter of *BdH-DG2-like*, one of the earliest leaf epidermal markers in *A. thaliana*^61^, helped to determine cells belonging to the epidermal tissue (Fig. 1E,G, Fig. S2L,M). Finally, we used homologs of *A. thaliana* cell cycle marker genes to assign cell cycle phases to the dataset (Fig. 1H). Four domains of mitotically active cells (i.e., G2-to-M phase cells) were identified; one domain lies between shoot apex and vasculature and expresses *BdWOX4* and *BdPIN1a*, one lies in the mesophyll and two in the epidermis (Fig. 1G,H, Fig. S1D,J).

Taken together, we generated a 70k-cells containing single-cell transcriptomic atlas of developing grass leaves from shoot apex to the mature leaf. A specific identity could be assigned to most clusters, dividing cells could be clearly identified and meristematic tissues could be distinguished from developing vasculature.

### Subsetting the epidermis identifies distinct epidermal lineages

We then subsetted the epidermal lineages and specifically reclustered and reanalyzed the developing grass leaf epidermis from protodermal stem cells to maturing and differentiating epidermal cell types (Fig. 2C, Fig. S3A,B). The mature abaxial leaf epidermis in *B. distachyon* consists of linear cell files that always have either stomatal complexes or prickle hair cells (HCs), spaced by at least one pavement cell (Fig. 2A). Above leaf veins, additional transverse divisions form silica cells below HCs. The grass leaf epidermis develops in a strictly linear manner, with seven well defined stages of stomatal development used as reference points throughout this manuscript (Fig. 2B).

**Figure 2.**
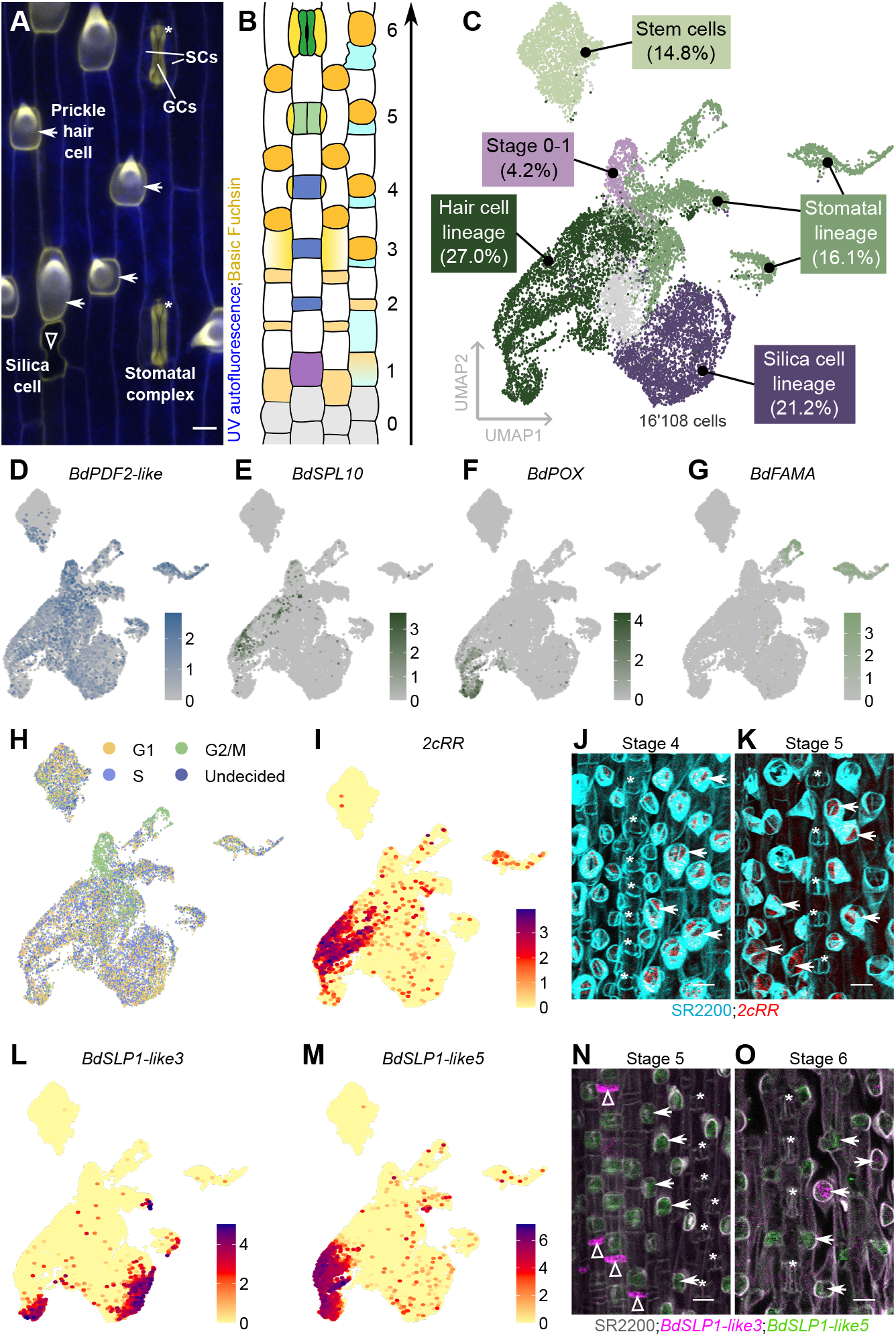
Subsetting the epidermis identifies epidermal lineages and novel marker genes. **(A)** Confocal image of the mature abaxial leaf epidermis of *B. distachyon*. Stomatal complexes are marked with asterisks, prickle hair cells with arrows and silica cells with triangles. Blue is cell wall autofluorescence and yellow is lignin stained by basic fuchsin (image adapted from ^43^). **(B)** Schematic of leaf epidermal developmental stages in *B. distachyon*. Protodermal stem cells (stage 0) acquire cell file identities (stage 1) and divide asymmetrically to form an apical guard mother cell (GMC) or hair cell (HC) lineage precursor and an interstomatal or inter-hair cell (stage 2). Inter-hair cells above leaf veins can divide again asymmetrically to form silica cell precursors. A GMC-derived signal initiates subsidiary mother cell (SMC) identity in lateral neighboring cells (stage 3). SMCs divide asymmetrically to form subsidiary cells (SCs) (stage 4). GMCs divide symmetrically to form two guard cells (GCs) (stage 5) which then differentiate into dumbbell-shaped GCs (stage 6). **(C)** Epidermis UMAP plot with epidermal cell type annotations (percentage of total cells in brackets). N = 16’108 cells. **(D)** Epidermis UMAP feature plot of epidermal marker *BdPDF2-like*. **(E)** Epidermis UMAP feature plot of HC marker *BdSPL10*. **(F)** Epidermis UMAP feature plot of HC marker *BdPOX*. **(G)** Epidermis UMAP feature plot of stomatal marker *BdFAMA*. **(H)** Epidermis UMAP plot with color indicating the cell cycle stage of each cell. **(I)** Epidermis UMAP feature plot of a two-component response regulator (*2cRR*). **(J, K)** Whole-mount RNA-FISH images of the *2cRR* (red) at developmental stages 4 (J) and 5 (K). Cell walls stained with SR2200 (cyan). Asterisks indicate stomatal complexes and arrows indicate HCs. **(L)** Epidermis UMAP feature plot of *BdSILIP-LANT1-like3* (*BdSLP1-like3*). **(M)** Epidermis UMAP feature plot of *BdSLP1-like5*. **(N, O)** Whole-mount multiplex RNA-FISH images of *BdSLP1-like3* (magenta) and *BdSLP1-like5* (green) at developmental stages 5 (N) and 6 (O). Cell walls stained with SR2200 (gray). Asterisks indicate stomatal complexes, arrows indicate HCs and triangles indicate silica cell precursors. Color legends in the UMAP feature plots show expression strength. Scale bars,10 µm. Also see Fig. S3.

We used epidermal marker genes to assign stomatal and HC lineage identities (Fig. 2C-G, Fig. S3D-M) and used cell cycle markers to assign cell cycle phases (Fig. 2H). Notably, there were two clusters with many mitotically active cells (Fig. 2H). The first one is likely the cells that divide during stage 0 and 1 of epidermal development without any clear indication of future cell fates. The second one is specific to the stomatal lineage and likely represents dividing GMCs (see below).

We then compared clusters of interest to all other epidermal clusters to identify differentially expressed, novel marker genes (DEGs) for different cell types (Table S4). Putative marker genes were chosen according to specific expression patterns in a given cell type or stage and were filtered to exclude protoplasting-affected genes (i.e., bulk RNA-sequencing data of non-protoplasted versus protoplasted tissues; (Fig. S3C, Table S5)). We then used whole-mount RNA fluorescence *in situ* hybridization (HCR™ RNA-FISH^62,63^) to verify the expression pattern of candidate marker genes *in planta*. A two-component response regulator (*2cRR*) was confirmed to be a new marker for HCs at stages 4 and 5 of epidermal development (Fig. 2I-K). For silica cells, the only known marker gene was the sorghum *SbSILIPLANT1* (*SbSLP1*), a gene suggested to precipitate silica in leaf silica cells^64,65^. Homologs of *SbSLP1* showed specific expression patterns (Fig. S3N-V); *BdSLP1-like1, 2, 3* and *4* showed strong *in silico* expression in silica cells and mature HC stages, while *BdSLP1-like5, 6, 7* and *9* seemed specific to HCs with *BdSLP1-like5* appearing during earlier stages already (Fig. 2L,M; Fig. S3N-V). We, therefore, chose *BdSLP1-like3* as a marker for silica cells and mature HCs and *BdSLP1-like5* as a marker for HCs only. Multiplexed RNA-FISH showed that at stage 5, *BdSLP1-like3* was expressed in silica cell precursors whereas *BdSLP1-like5* was exclusive to HCs (Fig. 2N). During stage 6, *BdSLP1-like3* stops to be expressed in silica cells and instead starts to be expressed in HCs, while *BdSLP1-like5* expression decreases (Fig. 2O).

In conclusion, subsetting the epidermal lineages successfully identified the HC and silica cell lineages, which were confirmed *in planta* using multiplexed whole-mount RNA-FISH of newly identified marker genes.

### Isolation of the stomatal lineage clusters identifies both grass stomatal cell types

Graminoid stomatal morphology is highly derived, with two dumbbell-shaped GCs that are flanked by lateral SCs (Fig. 2A,B). To decode the transcriptional dynamics of the developing grass stomatal lineages, we used known marker genes to isolate all stomatal lineage cells from the epidermal cluster (n = 4’675 cells, Fig. 2G, Fig. 3A, Fig. S3F-M, Fig. S4A,B). For example, the sequentially acting bHLH TFs *SPEECHLESS* (*SPCH*), *MUTE* and *FAMA*, were used to assign the GC lineage (Fig. 3C-G, Fig. S4D-H^9,44,54^;). The generation of a *BdFAMA* transcriptional reporter clearly demarcated the developmental stages expressing *BdFAMA* (Fig. 3H,I). Furthermore, cell cycle phase assignment once again revealed actively dividing cells at stage 0-1 and the symmetrically dividing GMCs (Fig. 3B), which also express *BdMUTE* and *BdFAMA* (Fig. 3D,G).

**Figure 3.**
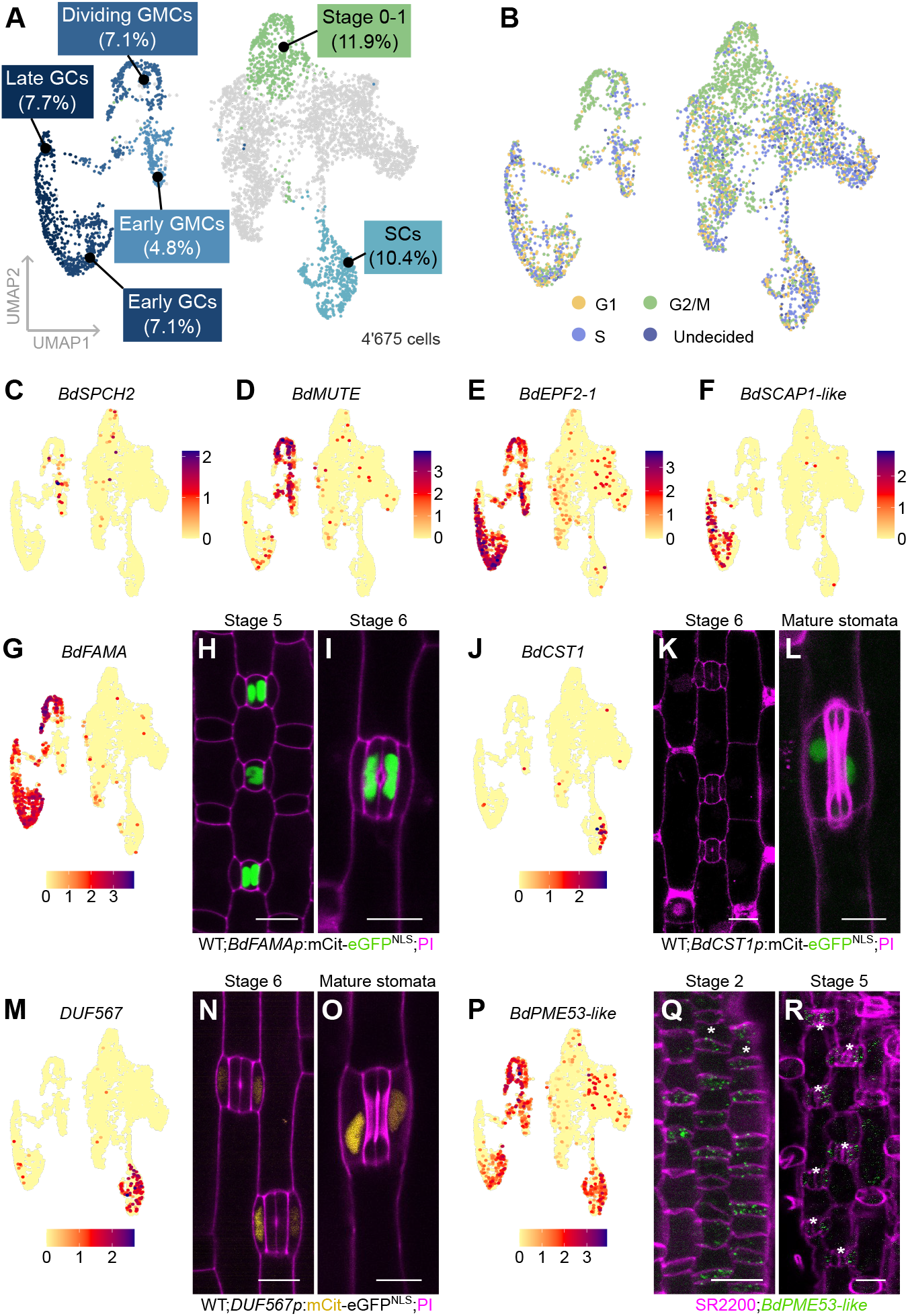
Spatiotemporal development of the stomatal lineage. **(A)** Stomatal lineage UMAP plot with stomatal cell types and stage annotations (percentage of total cells in brackets). N = 4’675 cells. **(B)** Stomatal lineage UMAP plot with color indicating the cell cycle stage of each cell. **(C)** Stomatal lineage UMAP feature plot of early stomatal lineage marker *BdSPCH2*. **(D)** Stomatal lineage UMAP feature plot of guard mother cell (GMC) marker *BdMUTE*. **(E)** Stomatal lineage UMAP feature plot of guard cell (GC) lineage marker *BdEPF2-1*. **(F)** Stomatal lineage UMAP feature plot of maturing GC marker *BdSCAP1-like*. **(G)** Stomatal lineage UMAP feature plot of GC lineage marker *BdFAMA*. **(H-I)** Expression of *BdFAMAp:mCitrine-eGFP*^*NLS*^ in developmental stages 5 (H) and 6 (I). **(J)** Stomatal lineage UMAP feature plot of mature subsidiary cells (SCs) marker *BdCST1*. **(K-L)** Expression of *BdCST1p:mCitrine-eGFP*^*NLS*^ in developmental stage 6 (K) and mature SCs (L). **(M)** Stomatal lineage UMAP feature plot of novel SC marker *DUF567*. **(N-O)** Expression of *DUF567p:mCitrine-eGFP*^*NLS*^ in developmental stage 6 (N) and mature SCs (O). Cell walls stained with propidium iodide (PI, magenta) in all reporter images. **(P)** Stomatal lineage UMAP feature plot of *BdPECTIN METHYLESTERASE 53-like* (*BdPME53-like*). **(Q-R)** Whole-mount RNA-FISH images of *BdPME53-like* (green) at developmental stages 2 (Q) and 5 (R). Cell walls stained with SR2200 (magenta). Asterisks indicate stomatal cell files (Q) or stomatal complexes (R). Color legends in the UMAP feature plots show expression strength. Scale bars, 10 µm. Also see Fig. S4.

The SCs were harder to identify as only a single, very late expressed SC-specific marker is known to date, *ZmCLOSED STOMATA 1* (*ZmCST1*; ^66^). In our first datasets we did not detect the expression of *BdCST1* and could therefore not rely on this gene to map SCs. To *bona fide* identify the SC population, two of the seven developmental zone libraries were produced from the *sid*/*bdmute-1* genotype that lacks SCs (Fig. S1C, Table S1; ^9^). Indeed, comparison of the two wild type (WT) libraries and the two SC-lacking *sid/bdmute-1* libraries processed and sequenced at the same time readily identified a cluster strongly enriched for WT cells (Fig. S4C). The most recent libraries contained many more SCs and–consequenctly–enabled the detection of *BdCST1* transcripts *in silico* (Fig. 3J). A *BdCST1* transcriptional reporter suggested that only very mature SCs express this marker (Fig. 3K,L). Differential gene expression analysis then identified novel stomatal marker genes. A *DUF567* gene showed strong *in silico* expression in the SC cluster (Fig. 3M). Within the epidermis, a transcriptional reporter for *DUF567* was indeed expressed very specifically in SCs from young stage 6 to mature stomatal complexes (Fig. 3N,O), earlier than *BdCST1*. Furthermore, the pectin methylesterase *BdPME53-like* was suggested to be expressed in the developing GC and SC lineages (Fig. 3P). Indeed, signal was detected from stage 2 GMCs to stage 5 GCs and in SCs from recruitment until maturing stomata (Fig. 3Q,R).

In conclusion, we successfully identified and confirmed the GC and SC lineages, found novel SC and GC markers, and cloned novel promoters specific to the SC lineage.

### Pseudotime trajectory of the grass guard cell lineage identifies stage-specific marker genes

Even though spatial (and spatiotemporal) information is lost in droplet-based scRNA-seq approaches, the temporal dynamics can be inferred using pseudotime analysis, particularly for lineages with sufficient *a priori* knowledge on developmental transitions like the GC lineage. Thus, we subsetted the GC lineage based on the stomatal bHLH TFs (n = 1’805 cells, Fig. 3C-G, Fig. 4A, Fig. S5A,B). Pseudotime analysis revealed the developmental gradient from early divisions at stage 0-1 towards fully differentiated GCs (Fig. 4A,B). Plotting the expression of known markers of different stages across the pseudotimeline confirmed the accuracy of the developmental GC trajectory (Fig. 4C).

**Figure 4.**
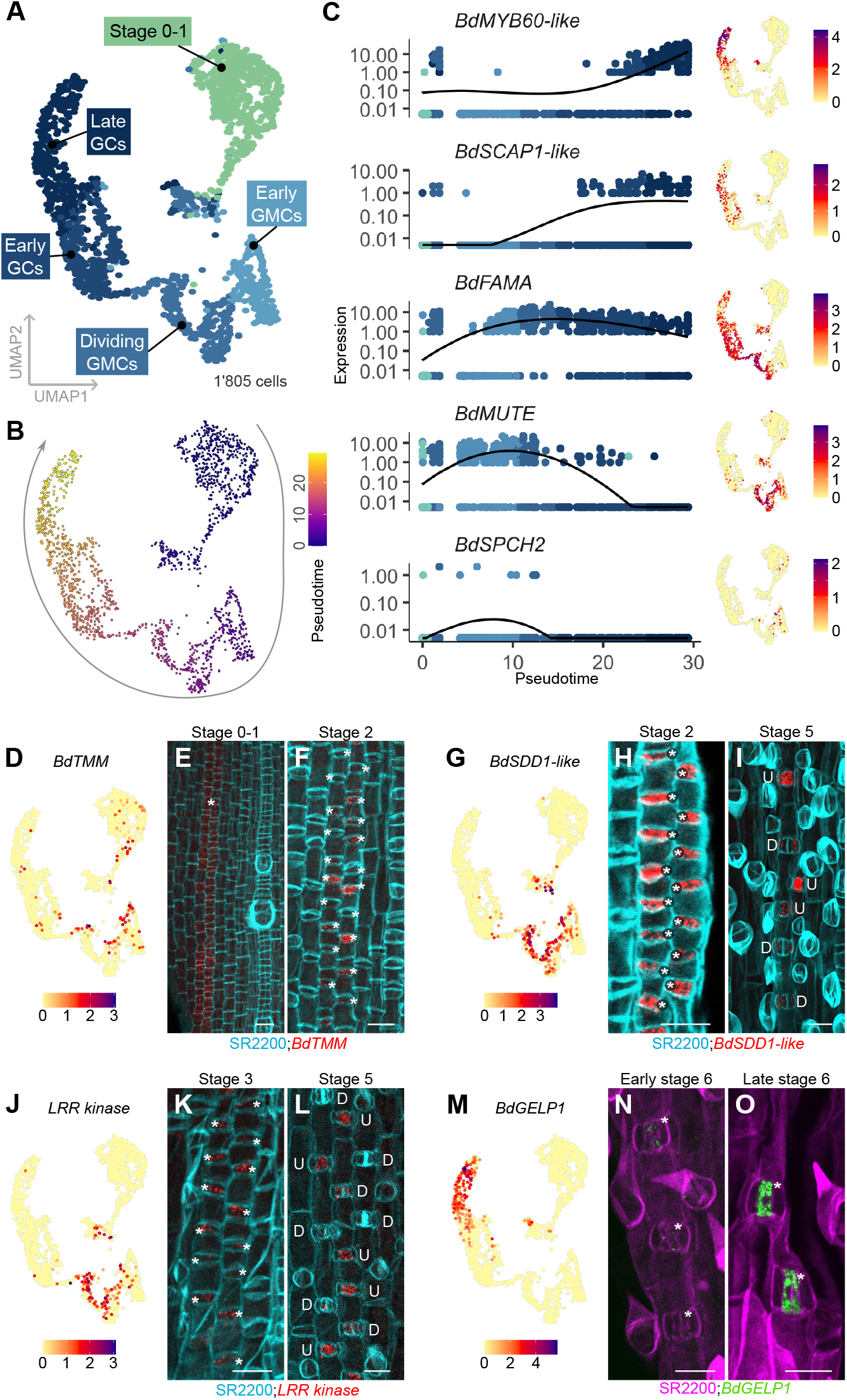
Sequential gene expression in the stomatal guard cell lineage. **(A)** UMAP plot with developing guard cell (GC) lineages; developmental stages are color-coded and annotated. N = 1’805 cells. **(B)** GC lineage UMAP plot with color indicating pseudotime. Arrow indicates developmental trajectory; heatmap indicates pseudotime stages. **(C)** Dot plot showing expression of marker genes along pseudotime. GC lineage UMAP feature plots of the respective genes are shown on the right. **(D)** GC lineage UMAP feature plot of *BdTOO MANY MOUTHS* (*BdTMM*). **(E, F)** Whole-mount RNA-FISH images of *BdTMM* (red) at developmental stages 0-1 (E) and 2 (F). Cell walls stained with SR2200 (cyan). Asterisks indicate stomatal cell files (E) and stomatal complexes (F). **(G)** GC lineage UMAP feature plot of *BdSTOMATAL DENSITY AND DISTRIBUTION 1-like* (*BdSDD1-like*). **(H**, Whole-mount RNA-FISH images of *BdSDD1like* (red) at developmental stages 2 (H) and 5 (I). Cell walls stained with SR2200 (cyan). Asterisks indicate guard mother cells (GMCs). “U” indicates undivided and “D” indicates divided GMCs. **(J)** GC lineage UMAP feature plot of *LEUCINE RICH REPEAT KINASE* (*LRR kinase*). **(K, L)** Whole-mount RNA-FISH images of *LRR kinase* (red) at developmental stages 3 (K) and 5 (L). Cell walls stained with SR2200 (cyan). Asterisks indicate guard mother cells (GMCs). “U” indicates undivided and “D” indicates divided GMCs. **(M)** GC lineage UMAP feature plot of *BdGDSL ESTERASE/LIPASE1* (*BdGELP1*). **(N, O)** Whole-mount RNA-FISH images of *BdGELP1* (green) at early (N) and late (O) developmental stage 6. Cell walls stained with SR2200 (magenta). Asterisks indicate stomatal complexes. Color legends in the UMAP feature plots show expression strength of the gene of interest. Scale bars, 10 µm. Also see Fig. S5.

We then selected several sequential and novel marker genes along this trajectory to confirm the developmental timeline from protodermal stem cells to maturing GCs *in planta*. The stomatal *BdTOO MANY MOUTHS* (*BdTMM*) co-receptor was suggested to be expressed from the earliest stages in the GC lineage (Fig. 4D, Fig. S5C). RNA-FISH revealed that *BdTMM* was truly expressed in very early, stage 0-1 stomatal files preceding the transverse asymmetric patterning division (Fig. 4E). After this division, the signal was mostly restricted to GMCs and disappeared after stage 2 (Fig. 4F). A homolog of the stomatal peptidase STOMATAL DENSITY AND DISTRIBUTION1 (SDD1; ^67^) showed strong *in silico* expression in young and dividing GMCs (Fig. 4G). Indeed, we found strong RNA-FISH signal for *BdSDD1-like* in stage 2 GMCs up until the symmetric division, after which it quickly disappeared, strongly matching the *in silico* expression prediction of our dataset (Fig. 4H,I). A *LEUCINE-RICH-REPEAT PROTEIN KINASE* (*LRR kinase*) showed a similar, yet slightly later *in silico* expression pattern compared to *BdSDD1-like* (Fig. 4J). Much like *BdSDD1-like*, signal of the *LRR kinase* probe was detected from stage 2 GMCs and abruptly disappeared in stage 5 as soon as the GMC had completed its division (Fig. 4K,L). Lastly, the *GDSL ESTERASE/LIPASE1* gene (*BdGELP1*) was suggested to be a mature GC marker gene (Fig. 4M), which was confirmed by RNA-FISH probes against *BdGELP1 in planta* (Fig. 4N,O).

We also performed pseudotime analysis for the HC lineage, to test if it can produce a useful trajectory with less *a priori* knowledge (Fig. S5D-F). Indeed, we found the expected sequential expression of the few currently known HC developmental marker genes (Fig. S5G) suggesting that our dataset paired with pseudotime analysis can reveal novel marker genes and developmental regulators of any epidermal lineage.

### Grass pavement cells are highly heterogenous

In contrast to stomatal cells, HCs or silica cells, no marker genes have been reported for pavement cells in grasses. We, therefore, used stomatal lineage-related genes to identify interstomatal cell clusters, which are stomatal lineage-derived pavement cells, and attempted to use these as a proxy to identify general pavement cell markers.

In *A. thaliana*, the ERECTA-YODA-MITOGEN-ACTIVATED PROTEIN (MAP) kinase (MAPK) signalling cascade ensures spacing of stomata by at least one pavement cell through the down-regulation of stomatal bHLH TFs upon reception of EPIDERMAL PATTERNING FACTOR 1/2 (EPF1/2) peptide ligands that are secreted by developing stomatal complexes (reviewed in ^68,69^). In grasses, the inhibitory role of the pathway in epidermal cell file patterning seems to be conserved. For example, EPF ligands were shown to affect stomatal density (e.g. ^70–72^) and *yoda1* mutants in *B. distachyon* and barley caused within-file clustering of specialized cells^73,74^.

We hypothesized that the ERECTA-YODA-MAPK signalling pathway is preferably active in interstomatal pavement cells and analyzed the *in silico* expression patterns of *B. distachyon* homologs of the inhibitory signalling pathway. As expected, *BdEPF2-1* and *BdEPF2-2* ligands are primarily expressed in the GC lineage (Fig. 3E, Fig. S4H). *BdYDA1/2* and the putative downstream MAPKs *BdMKK4/5-like1, BdMKK4/5-like2, BdMPK3-like* and *BdMPK6-like* were expressed rather ubiquitously, yet a slight enrichment in the right-most clusters was observed (clusters 1, 3, and 8; Fig. S4B,I-N). Similarly, *BdERECTA, BdERECTA-LIKE1* (ERL1) and three co-receptors *SOMATIC EMBRYOGENESIS RECEPTOR KINASE* (*BdSERKs*) were strongly and preferentially expressed in clusters 1, 3 and 8 (Fig. 5A-E, Fig. S4B), suggesting that these not yet annotated cells could potentially be the interstomatal cells (highlighted cells, Fig. 5N). Indeed, RNA-FISH against *BdERL1* confirmed its expression in stomatal cell files from very early stages of stomatal development through to GMCs and interstomatal cells after the transverse patterning division at stage 2 (Fig. 5F-H).

**Figure 5.**
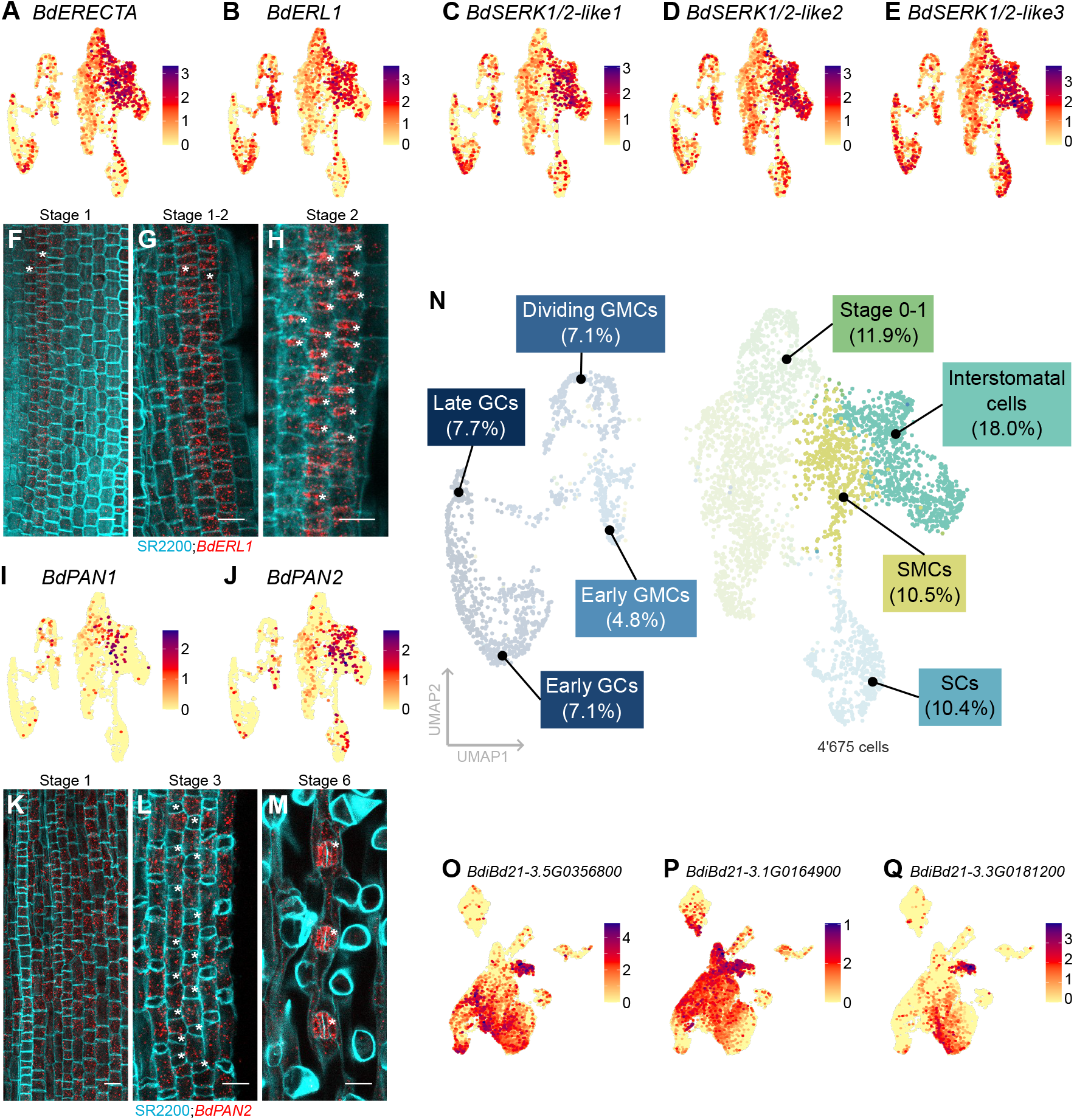
Analysis of interstomatal pavement cells as a proxy to identify transcriptomic pavement cell features. **(A-E)** Stomatal lineage UMAP feature plot of *BdERECTA* (A), *BdERECTA-LIKE1* (*BdERL1*) (B), *BdSOMATIC EMBRYOGENESIS RECEPTOR KINASE* (*BdSERK1/2-like1*) (C), *BdSERK1/2-like2* (D), and *BdSERK1/2-like3* (E). **(F-H)** Whole-mount RNA-FISH images of *BdERL1* (red) at developmental stage 1 (F), stage 1 to 2 (G) and stage 2 (H). Cell walls stained with SR2200 (cyan). Asterisks indicate stomatal cell files (F, G) and guard mother cells (GMCs, H). **(I-J)** Stomatal lineage UMAP feature plot of *BdPANGLOSS1* (*BdPAN1*) (I) and *BdPAN2* (J). **(K-M)** Whole-mount RNA-FISH images of *BdPAN2* (red) at developmental stage 1 (K), 3 (L) and 6 (M). Cell walls stained with SR2200 (cyan). Asterisks indicate GMCs (L) and stomatal complexes (M). **(N)** Stomatal lineage UMAP plot with stomatal cell type and stage annotations (percentage of total cells in brackets). Same plot as in Fig. 3A but with interstomatal cells and subsidiary mother cells (SMCs) labelled. **(O-Q)** Epidermis UMAP feature plots of marker genes of the interstomatal cells clusters. Color legends in the UMAP feature plots indicate expression strength. Scale bars, 10 µm. Also see Fig. S4.

Such interstomatal cells can also be stomatal SMCs when two adjacent stomatal rows are formed. However, no exclusive SMC markers are known. The main SMC identity factor *BdMUTE* is not expressed in SMCs but rather translocates from GMCs to SMCs^9,53^ and many of the genes affecting SMC polarization and division are not specific to SMCs (Fig. S4O-T; ^43^). Nonetheless, the two polarization factors *BdPANGLOSS1* (*BdPAN1*) and *BdPAN2*^52,75,76^ appeared to be quite strongly expressed in the putative interstomatal cells (Fig. 5I,J). *BdPAN1* showed strongest ^77^ expression in cluster 3, suggesting that this cluster corresponded to SMCs (Fig. 5I,N, Fig. S4B). RNA-FISH confirmed *BdPAN2* expression in interstomatal cells, SMCs, and pavement cells in hair cell files (Fig. 5K,L). Beyond stage 2 and 3, *BdPAN2* seemed to be strongly expressed in SCs and, later, in GCs, a pattern that matched its described function in SC shape and *in silico* expression in our atlas (Fig. 5M). Together, this suggested that clusters 1, 3, and 8 are interstomatal cells, with cluster 3 potentially being enriched for SMCs (Fig. 5N, Fig. S4B).

We then attempted to identify marker genes in clusters 1, 3 and 8 that might be general pavement cell markers. When mapping some of these putative pavement cell markers onto the epidermal dataset, we got either very broad, non-specific expression patterns (Fig. 5O, Fig. 2C) or markers for what appears to be developmentally younger cells (Fig. 5P, Fig. 2C) or marker genes that are expressed alongside the hair or silica cell domain in addition to interstomatal cells (Fig. 5Q, Fig. 2C).

Together, this indicated that pavement cells are highly heterogenous and their transcriptional identity might vary significantly depending on whether they stem from the stomatal, the hair cell or the silica cell lineage. This might explain why pavement cells do not form specific, readily identifiable lineage clusters in grasses.

### Gene regulatory network analysis identifies key transcription factors as main regulons in the stomatal lineage

To identify the main TF hubs and their putative target genes in the stomatal clusters, we conducted gene regulatory network (GRN) analysis of the stomatal subset using MINI-EX (version 3; ^78,79^). The GRN analysis revealed different TF regulons that may be key to regulating the developmental progression of cell types at different developmental stages. Among the top five most relevant TFs per each stomatal lineage cluster, we found several that were already implicated in leaf and stomatal development (Fig. 6A, Fig. S6A, Table S6).

**Figure 6.**
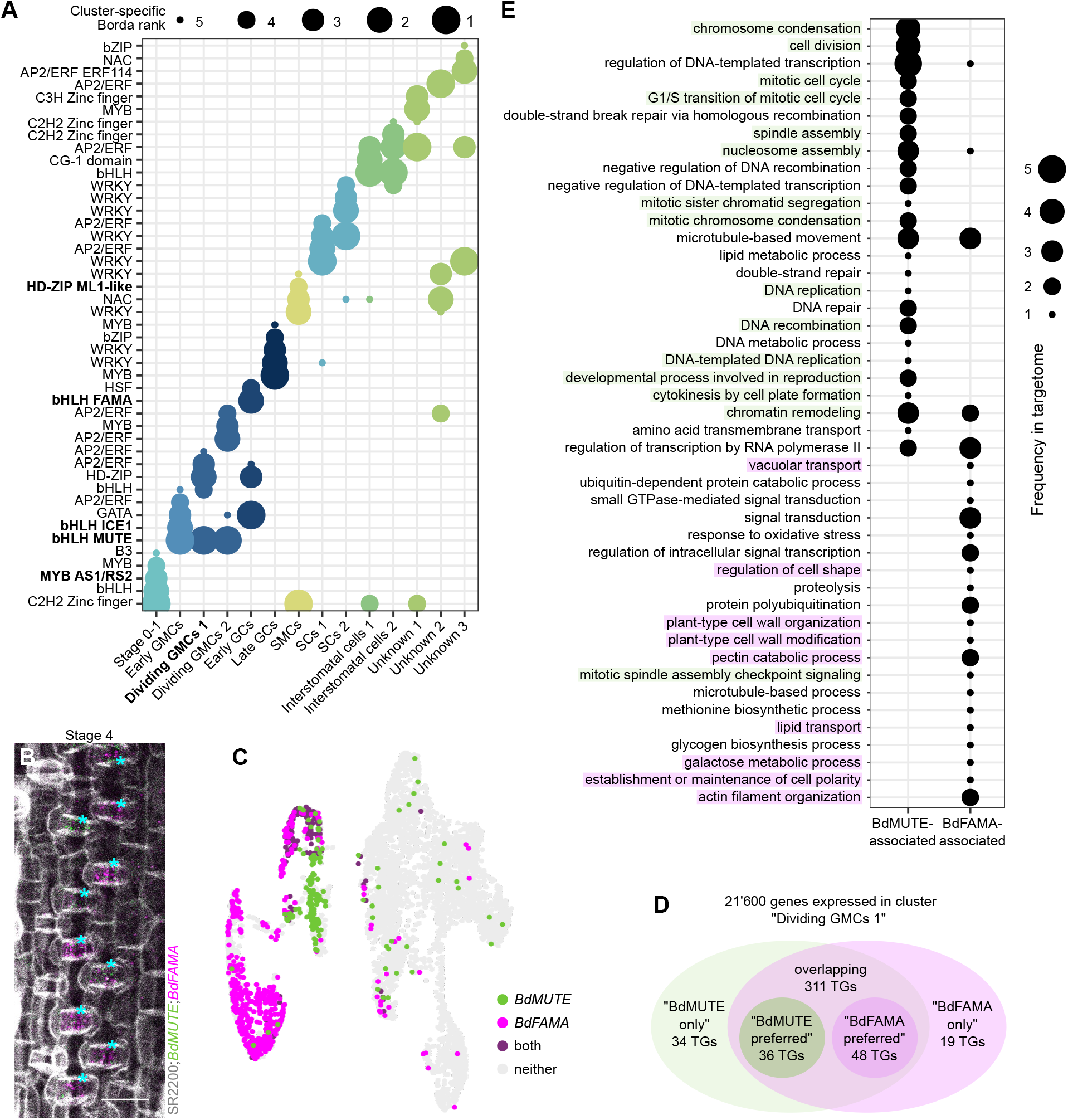
Gene regulatory network analysis of the stomatal lineages. **(A)** Dot plot showing the top 5 regulons (= transcription factors (TFs)) per cluster. TFs highlighted in bold have already been described in the context of leaf development. **(B)** Whole-mount multiplex RNA-FISH image of *BdMUTE* (green) and *BdFAMA* (magenta) at developmental stage 4. Cell walls stained with SR2200 (gray). Asterisks indicate stomatal complexes. Scale bar, 10 µm. **(C)** Stomatal lineage UMAP feature plot of *BdMUTE* (green) and *BdFAMA* (magenta). Cells expressing both genes are colored dark purple, cells expressing neither are in gray. **(D)** Venn diagram of target genes (TGs) of BdMUTE (green) and BdFAMA (magenta) in the “Dividing GMCs 1” cluster. 34 TGs are exclusive to BdMUTE (“BdMUTE only”), 19 TGs are exclusive to BdFAMA (“BdFAMA only”) and 311 genes are targeted by both TFs. Of these, 36 show preferential co-expression with BdMUTE (logarithmic weight ratio >3) and 48 show preferential co-expression with BdFAMA (logarithmic weight ratio <3). **(E)** Dot plot showing the frequency of the top 25 GO terms for “BdMUTE-associated” TGs (“BdMUTE only” and “BdMUTE preferred”) and “BdFAMA-associated” TGs (“BdFAMA only” and “BdFAMA preferred”). GO terms associated with cell division are marked in green and GO terms associated with cell morphogenesis are marked in magenta. Also see Fig. S6.

Within the early cluster of stage 0-1 we found *BdASYMMETRIC LEAVES 1* (*BdAS1*), a TF involved in proximo-distal patterning of early leaves^80^, in third position (Fig. 6A, Fig. S6A,B, Table S6). *BdPROTODERMAL PATTERNING FACTOR 2-like* (*BdPDF2-like*), which is closely related to *AtPDF2* and defines L1/epidermal fate in the *thaliana* shoot apex^81^, was in sixth position (Fig. S6C, Table S6). Finally, a homolog of the leaf morphogenesis TF *AtTEOSINTE BRANCHED 1, CYCLOIDEA, PROLIFERATING CELL FACTOR1/2 4* (*AtTCP4*; ^82^), *BdTCP4-like*, was found in seventh position (Fig. S6D, Table S6).

*BdMUTE* is the top regulon in the three GMC clusters in line with its role in regulating GMC cell division orientation (Fig. 6A; ^53^). *BdICE1* is in second position for the early GMCs suggesting it to be the putative heterodimerization partner of *BdMUTE* (Fig. 6A; ^54^). *BdTSO1*, whose *A. thaliana* ortholog was found to be involved in the GMC to GC transition^83^, is in eighth position at this stage (Fig. S6E, Table S6). *BdFAMA*, shown to regulate GC differentiation and pore formation^54^, was the second most relevant regulon in early GCs (Fig. 6A, Table S6) and its putative heterodimerization partner *BdSCRM2* was in seventh position in early GCs (Table S6; ^44,54^).

In conclusion, single-cell GRN analysis revealed core TF hubs in accordance with their implied function from molecular and genetic analysis during grass stomatal development and provides a valuable resource to find novel key regulators in the developmental stages of interest.

### Distinct targetomes of BdMUTE and BdFAMA in guard mother cells

The MINI-EX output also yields target genes (TGs) per regulon (targetomes), if known TF motifs and a list of all genes that contain such motifs in their cis-regulatory regions (generated with the FIMO tool^84^), are used as input. This enabled cluster-specific analyses of putative downstream genetic programs for each identified regulon per cluster.

The two bHLHs *BdMUTE* and *BdFAMA* overlap in their expression windows in GMCs both *in silico* and *in planta* (Fig. 6B,C), yet have distinct functions in regulating division orientation and GC differentiation, respectively^53,54^. Therefore, they provide an excellent example to evaluate putative differences in the targetomes of two closely related TFs with the same binding motif (CANNTG bHLH motif; ^85,86^) yet distinct functional roles. In the cluster “Dividing GMCs 1” (cluster 7, Fig. 6A, Fig. S4B), 21’600 genes are expressed, of which 311 appeared as TGs for both TFs, 34 were exclusive to *BdMUTE* and 19 were exclusive to *BdFAMA* (Fig. 6D, Table S7). Each TF-TG combination was assigned a weight value describing the strength of co-expression between the two genes. To identify preferential TGs, we calculated the ratio of weight_BdMUTE_ by weight_BdFAMA_ and filtered for TGs that had a logarithmic weight ratio of higher than 3 to obtain genes that have stronger association with *BdMUTE* and lower than −3 for genes that are more linked to *BdFAMA*. This resulted in additional 36 preferential TGs for *BdMUTE* and 48 preferential TGs for *BdFAMA* (Fig. 6D). We then explored the functional annotations of exclusive and preferential TGs using gene ontology (GO) terms (Fig. 6D,E). Notably, the *BdMUTE* TGs are more likely to be involved in mitotic processes (Fig. 6E, green) whereas the *BdFAMA* targetome seemed to be rather associated with cell wall biology and cellular morphogenesis (Fig. 6E, magenta), consistent with their genetically determined functions. The functionally divergent TG-set enrichment of *BdMUTE* and *BdFAMA* suggested that our atlas allows for the discovery of cell-type and TF-specific gene networks in a highly targeted manner.

### From division control to stomatal density-functional analysis of two developmental marker genes

We selected two marker genes for mutant analysis to show how this dataset can help to identify new players in leaf or stomatal development. First, a GRAS TF, *BdGRAS32*, which is the orthologue of the rice TF *DWARF AND LOW-TILLERING (DLT*)/*OsGRAS-32*^87^, was identified as a putative cell division factor throughout leaf development as its *in silico* expression strongly overlapped with actively dividing cells (Fig. 7A,B). RNA-FISH verified that the gene was already expressed in very young leaf primordia at the shoot apex and in the earliest stages of epidermal development (Fig. 7C,D). Finally, strong expression could also be detected in GMCs shortly before the division at stage 5 (Fig. 7E). This suggested a role in regulating cell division during *B. distachyon* leaf development. To investigate this, we generated a *bdgras32* CRISPR mutant (Fig. 7F). We detected misdivisions with seemingly misplaced, additional division planes around GCs and in SCs (Fig. 7G). In addition, imaging and segmentation of the developing leaf epidermis (stage 1-2) of wild type and *bdgras32* mutants (Fig. 7I) showed significantly higher cell numbers and, as a consequence, smaller cells in *bdgras32* compared to wild type (Fig. 7J,K). Together, we propose that *BdGRAS32* represses cell division capacity in the developing *B. distachyon* leaf.

**Figure 7.**
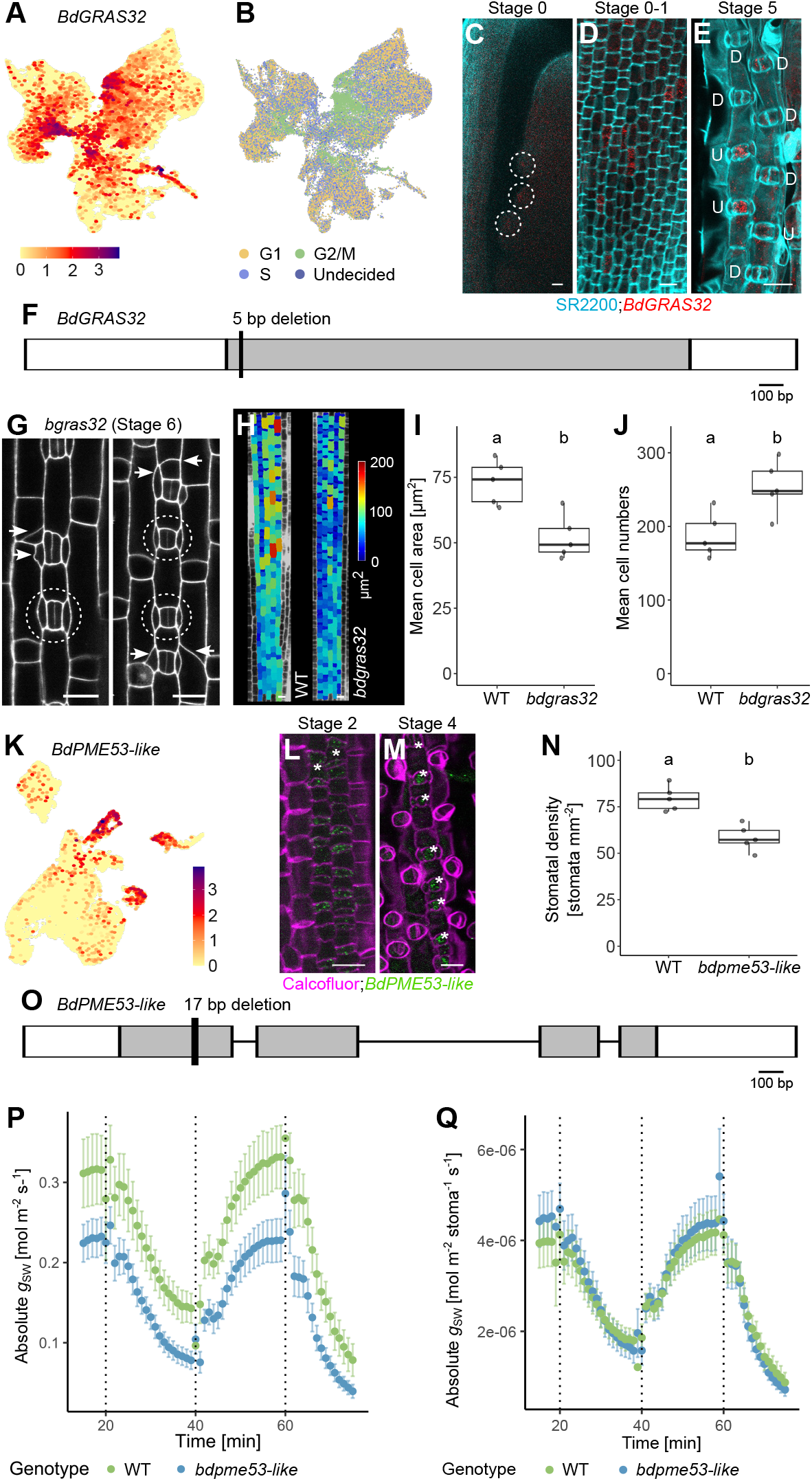
Functional analysis of selected marker genes. **(A)** Whole dataset UMAP feature plot of *BdGRAS32*. **(B)** Whole dataset UMAP plot with color indicating the cell cycle stage of each cell. Same plot as in Fig. 1H. **(C-E)** Whole-mount RNA-FISH images of *BdGRAS32* (red) in leaf primordia (C, circled), epidermis developmental stage 0-1 (D) and 5 (E). Cell walls stained with SR2200 (cyan). “U” indicates undivided and “D” indicates divided guard mother cells (GMCs). Scale bars, 10 µm. **(F)** Gene model of *BdGRAS32* with the mutation of the *bdgras32* mutant indicated. **(G)** Mutant phenotype of *bdgras32* at developmental stage 6. Arrows indicate misdivisions, ellipses indicate wild type (WT)-like stomatal complexes. Scale bars, 10 µm. **(H)** Representative MorphoGraphX heat map images of WT and *bdgras32*. Color indicates cell area. Scale bar, 50 µm. **(I)** Mean cell area of WT and *bdgras32*; n = 5 individuals per genotype. **(J)** Mean cell numbers of WT and *bdgras32*; n = 5 individuals per genotype. **(K)** Epidermis UMAP feature plot of *BdPECTIN METHYLESTERASE 53-like* (*BdPME53-like*). **(L, M)** Whole-mount RNA-FISH images of *BdPME53-like* (green) in developmental stage 2 (M) and 4 (N). Cell walls stained with Calcofluor (magenta). Asterisks indicate stomatal cell files (M) and stomatal complexes (N). Scale bars, 10 µm. **(N)** Stomatal density of WT and *bdpme53-like*; n = 5 individuals per genotype. **(O)** Gene model of *BdPME53-like* with the mutation of the *bdpme53-like* mutant indicated. **(P, Q)** Absolute stomatal conductance (*g*_SW_) of WT (green) and *bdpme53-like* (blue) per leaf area (Q) and per stoma (R). Standard error is indicated by error bars. Color legends in the UMAP feature plots indicate expression strength. Significant differences calculated with unpaired, two-sided Student’s *t*-test (p<0.5) are indicated by differing letters.

To functionally describe a stomatal marker gene, we chose to gene edit *BdPME53-like* (Fig. 7L,P), which we established as a marker for the developing GC and SC lineage (Fig. 3P-R, Fig. 7M,N). We hypothesized that mutating *BdPME53-like* could affect cell wall properties and infer with biomechanical aspects of stomatal opening and closing. Indeed, leaf-level gas exchange measurements indicated a decrease in stomatal opening, represented by lower stomatal conductance to water vapour (*g*_SW_, Fig. 7Q). Analysis of stomatal anatomy, however, showed that *bdpme53-like* formed significantly fewer stomata (Fig. 7O). By correcting the gas exchange values for averaged stomatal density per genotype, *g*_SW_ was no longer lower in the mutant on a per-stoma basis (Fig. 7R), suggesting that the observed physiological phenotype might not be due to impaired stomatal function but a decrease in stomatal density in *bdpme53-like*. In conclusion, our proof-of-concept functional validation of two genes strongly suggested that the *B. distachyon* single-cell leaf atlas can be leveraged to identify developmental factors that shape the functional anatomy of the most important photosynthetic powerhouse for ecosystems and humanity alike.

### An online gene expression viewer for the community

To make this dataset readily accessible, we created an expression browser tool to visualize expression of any *distachyon* gene of interest in our leaf atlas (Fig. S7G). We generated different tabs so that gene expression can be visualized with feature plots of the whole dataset, the epidermal subset or the stomatal subset (https://shiny.ips.unibe.ch/). Next to the feature plot, the expression viewer displays a UMAP plot with annotated tissues for reference. In addition, we computed a ggPlantmap^88^ visualization of the *bona fide* assigned tissues and cell types for an eFP-browser-like experience. A download function generates an overview of all tissues, the epidermal and the stomatal subset (feature plot of the gene and UMAP plot with tissue identities) and the ggPlantmap overview (Fig. S7G). We hope that this “*B. distachyon* leaf gene expression viewer” will be used by the research groups working primarily on *B. distachyon* and the extended grass developmental research community.

## Discussion

The genetic programs that pattern and coordinate the development of different tissues and cell types in grass leaves are still largely unexplored. In the past, gene discovery mainly relied on mutant screens or analysis of homologs of genes involved in leaf development in *A. thaliana*. The advent of single-cell/nucleus transcriptomics nowadays facilitates gene discovery, gene network and gene co-expression analysis in an unbiased, genome-wide and model system-independent manner^14^.

Here, we provide a single-cell transcriptomic leaf atlas of the wild model grass *B. distachyon* that comprises almost 70k cells from vegetative shoot apex to leaf developmental zone (Fig. 1). We used known markers of different tissues, cell types and stages to spatially and pseudotemporally resolve cluster identities (Fig. 1-5). While this study focused mostly on the leaf epidermis, we were also able to identify distinct expression patterns of developmental markers within the shoot apex and vasculature (Fig. S1-2). We predict that in-depth analysis of the vascular and mesophyllar subsets will allow the discovery of new developmental regulators along the spatiotemporal trajectories of these tissues even though some of the very large cells or tissues with massive secondary cell wall deposition and modification might not have been quantitatively captured.

Furthermore, we were not able to identify one rather enigmatic leaf epidermal cell type, the bulliform cells. We hypothesize that this is caused by their large cell size and/or physiological properties. Bulliform cells are rather long and bulging cells of the adaxial epidermis and there is a high probability that they were size filtered as the 10X Genomics platform has a cell size limit of around 40 µm^14^. In addition, bulliform cells are functionally associated with leaf rolling and water release under arid conditions^89,90^. Therefore, these cells might experience rather high cell turgor under well-watered conditions and are thus likely to burst when cell walls are digested to release protoplasts. A different sequencing platform designed to accommodate bigger cells (e.g. BD Rhapsody) or the use of nuclei rather than protoplasts can overcome cell size restrictions and protoplast bursting in future. Similarly, mature pavement cells might also be missing due to size exclusion, bursting or suboptimal protoplasting. The strong presence of *ER, ERL1* and *SERKs* in certain clusters of the epidermis, however, indicated that we were able to obtain at least younger, and therefore smaller, interstomatal pavement cells (Fig. 5). However, we were not able to find pavement cell-specific markers. We hypothesize that pavement cell transcriptomes in the strictly linearly patterned grass leaf epidermis tend to resemble their respective sister cells within a cell file, due to their shared origin. Therefore, we expect the young pavement cells to cluster close to their specialized neighbor cells, as appears to be the case for the interstomatal cells (Fig. 5).

Next, pseudotime analysis nicely recapitulated the developmental transitions of the GC lineage and HC lineage. For both lineages, however, stage-specific marker genes were known before, in particular for the GC lineage, where many of the early core developmental regulators are identified and genetically characterized^44,53,54^. This suggested pseudotime analysis to be a very powerful tool. Yet, such analysis will also forcibly yield a trajectory that might be biologically non-sensical. Some *a priori* knowledge of the biological context is required and biologically meaningful, ontogenetically related cell lineages should be isolated prior to pseudotime analysis.

We then implemented single-cell-resolved GRN analysis (using MINI-EX), which enabled the investigation of the targetomes of selected TFs within or across selected clusters. The comparison of two closely related TFs, *BdMUTE* and *BdFAMA*, which are both expressed in the dividing GMCs but regulate GMC division and GC differentiation, respectively^53,54^, showed that we are able to distinguish these two biological processes within a selected cluster based on the targetomes of the two distinct regulons (Fig. 6). Importantly, this method will allow comparative approaches between different sc-RNA-seq datasets of closely and distantly related species that use the same TFs for distinct stomatal morphologies and compositions (i.e. grasses and eudicots) as the targetome is more likely to be modified by evolution than the regulon (i.e., TF) itself^91–93^. Despite its elegance and power, there are limitations to the GRN analysis applied here. Firstly, it relies on transcriptional expression of a TF in the cell type of interest. This suffices for the regulatory analysis within that cell type, but mobile, non-cell autonomously acting TFs are excluded. *BdMUTE*, for example, is transcriptionally expressed and has a functional role in the GMCs, but its primary role is moving from GMCs to neighboring cells and establishing the SC lineage^9,53^. Our GRN approach can therefore only decipher putative targets of BdMUTE within the GMCs, but not in the SMCs. In addition, the targetome analysis is based on genes being co-expressed with a regulon and will therefore only yield target genes that are positively regulated by the regulon but might miss downregulated target genes.

To demonstrate how our dataset can be used to find new components of leaf development, we selected two genes with a specific expression pattern for functional analysis. Mutations in *BdGRAS32*, which is specifically expressed in mitotically active cells, showed smaller, more numerous cells and division defects (Fig. 7G-K). This confirms a similar phenotype found in rice mutants, where *dlt/osgras-32* mutants showed increased leaf width and higher cell numbers by regulating cyclin-related genes^87^. It is therefore likely that *BdGRAS32* also restricts cell division through regulation of cell cycle genes in *B. distachyon*. Mutations in the stomata-specific gene *BdPME53-like* affected stomatal density and, consequently, leaf-level gas exchange (Fig. 7L-R). This suggests a role of *BdPME53-like*-mediated cell wall modifications during stomatal development which may influence stomatal lineage patterning in the epidermis. In *A. thaliana*, the mutant of the closest related gene, *AtPME53*, had slightly increased stomatal length, width, and stomatal density, yet decreased stomatal aperture^94^. As we observed an opposing phenotype in regards to stomatal density, more work will be required to determine how *AtPME53* and *BdPME53-like* influence stomatal lineage development and why this may lead to opposite phenotypes.

In conclusion, we present a spatio-pseudotemporally resolved single-cell atlas of a developing grass leaf from leaf primordia at the shoot apex to mature leaf tissues in the wild grass model *B. distachyon*. This dataset can be used to identify and select genes in a targeted manner for downstream functional analyses to further decode leaf developmental processes in the most important plant family for humanity–the grasses.

## Supplemental Information

Document S1. Figures S1–S6, Tables S1-S3, and supplemental references

Table S4.

Table S5.

Table S6.

Table S7.

## Acknowledgments

We would like to thank the research gardeners Michael Schilbach (COS Heidelberg, Germany), Sarah Dolder, Jasmin Sekulovski and Christopher Ball (all University of Bern, Switzerland). For sequencing support, we acknowledge the Deep Sequencing Core Facility at Heidelberg University (D. Ibberson) and the Next Generation Sequencing Platform at the University of Bern (D. Steiner, P. Nicholson). Microscopy was performed on equipment supported by the Microscopy Imaging Center (MIC) at the University of Bern. Thank you to Ole-Tobias Schwarz, Alexander Kashev and the UBELIX HPC cluster at University of Bern for informatics support. Our thanks also goes to the Cereal Stem Cell Systems (CSCS) research unit for feedback and discussions, and Thorsten Schnurbusch (IPK Gatersleben) for providing the grass WOX phylogeny. This work was supported by the DFG FOR5235 “Cereal Stem Cell Systems” and the University of Bern.

## Author Contributions

Conceptualization: MTR; Data curation: LSB, IHP; Formal analysis: LSB, IHP, AG; Funding acquisition: RS, MTR; Investigation: LSB, PRD, IHP, NL, HL, IV, JEM, RPS; Methodology: LSB, PRD, IHP, RT, HL, IV, JEM, RS, MTR; Project administration: MTR; Resources: RS, MTR; Supervision: RS, MTR; Visualization: LSB, MTR; Writing—original draft: LSB, MTR; Writing—review & editing: all authors. All authors have read and approved the final manuscript.

## Materials & Methods

Key resources table

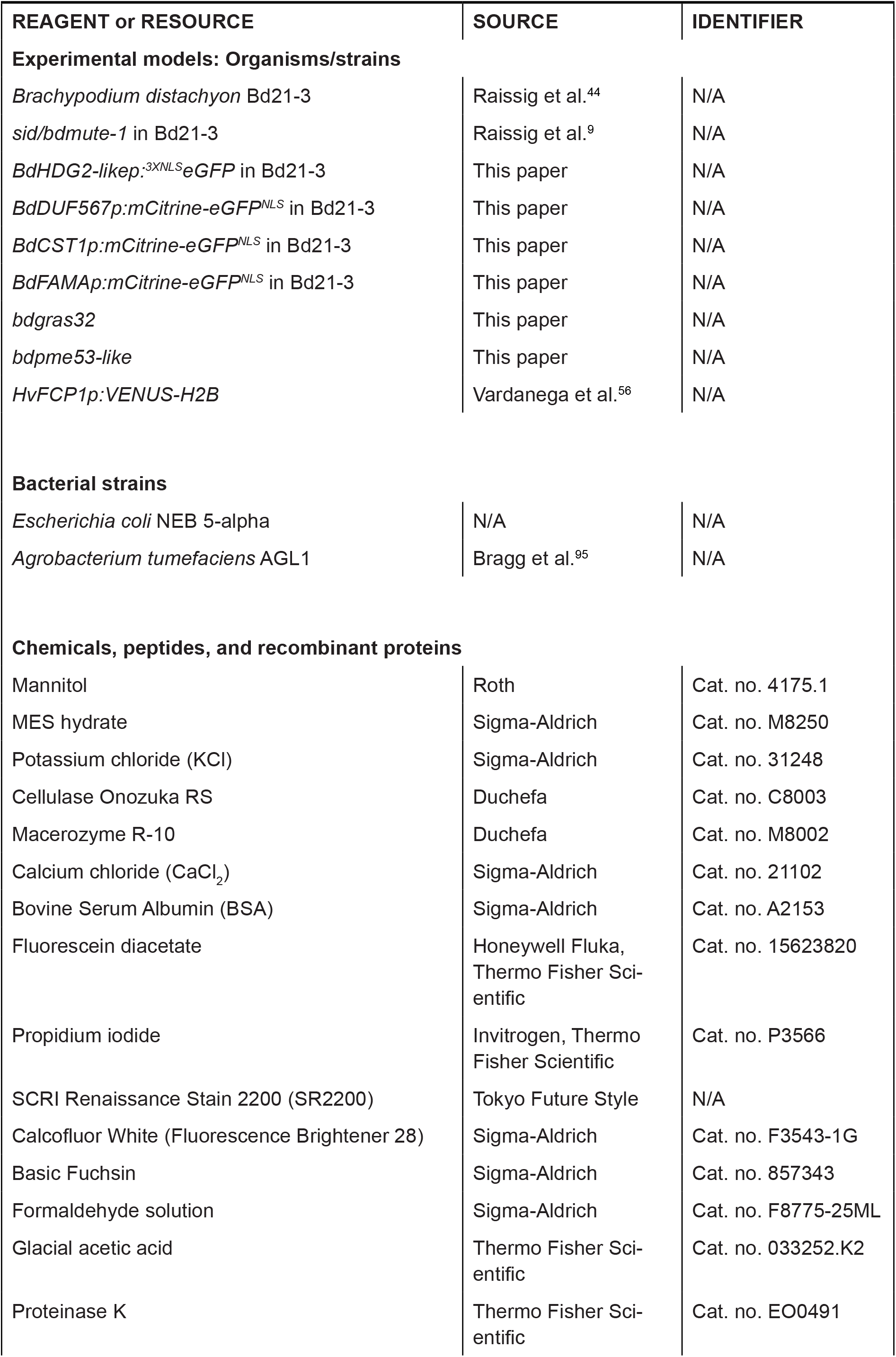
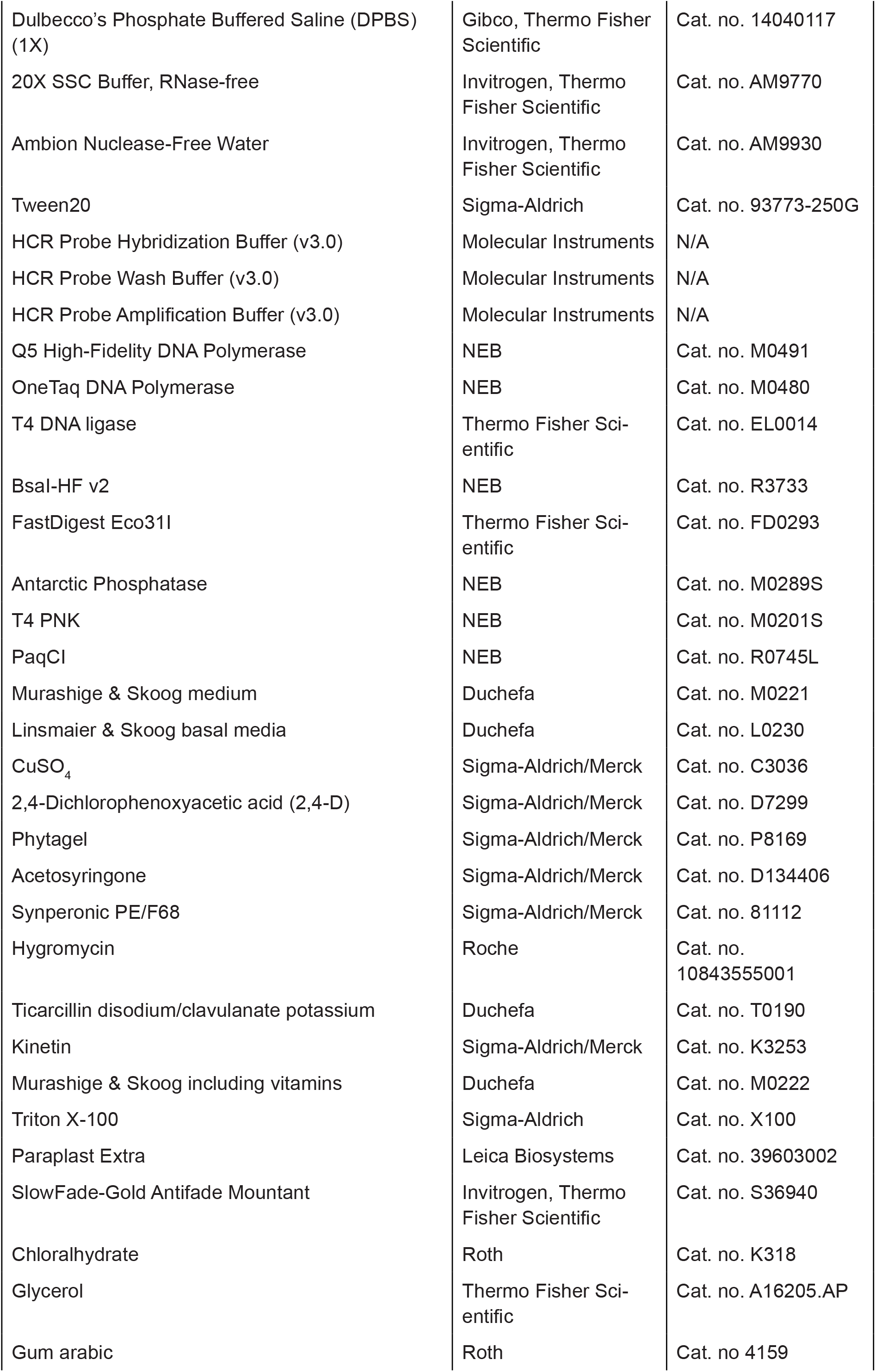
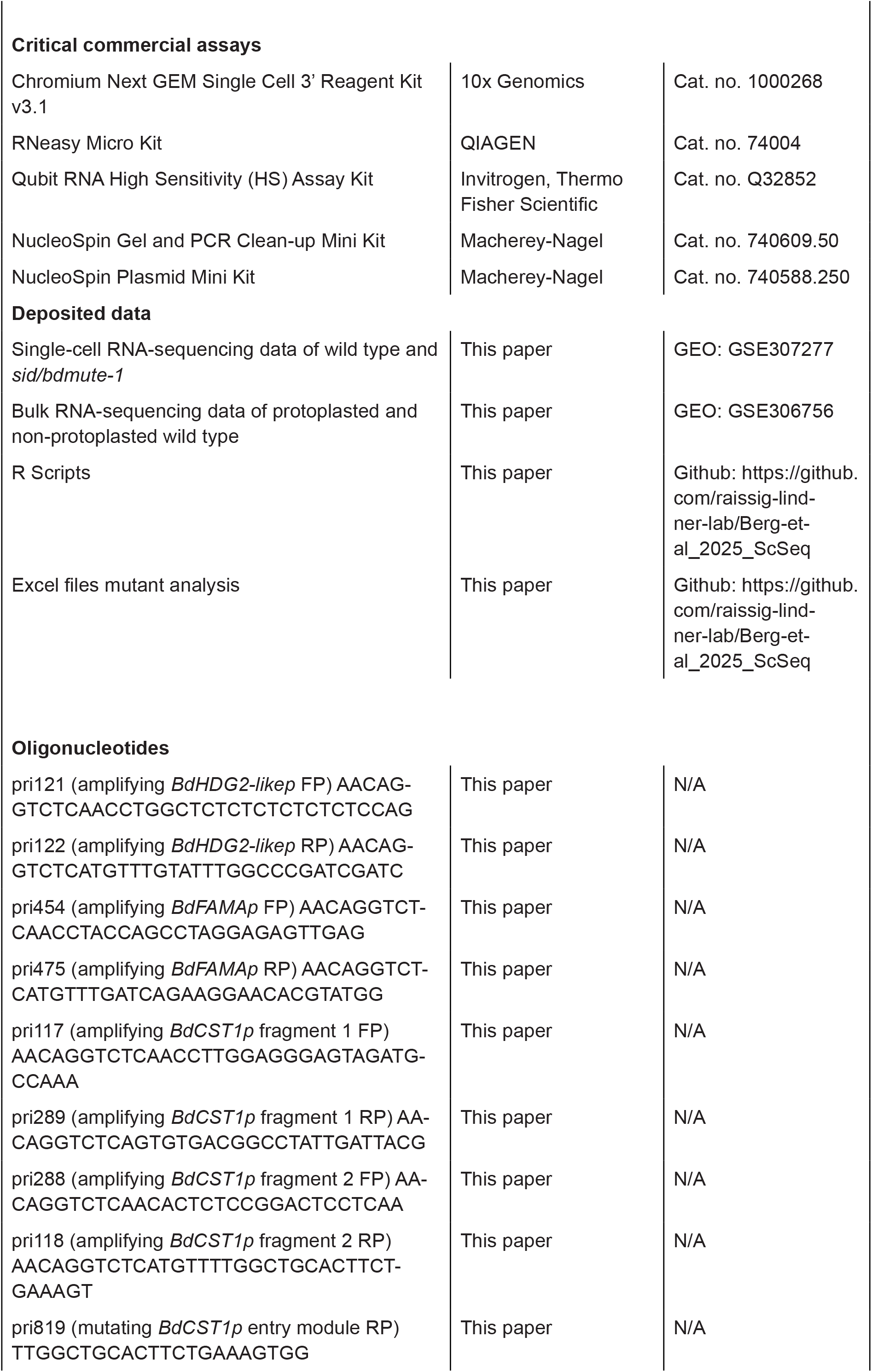
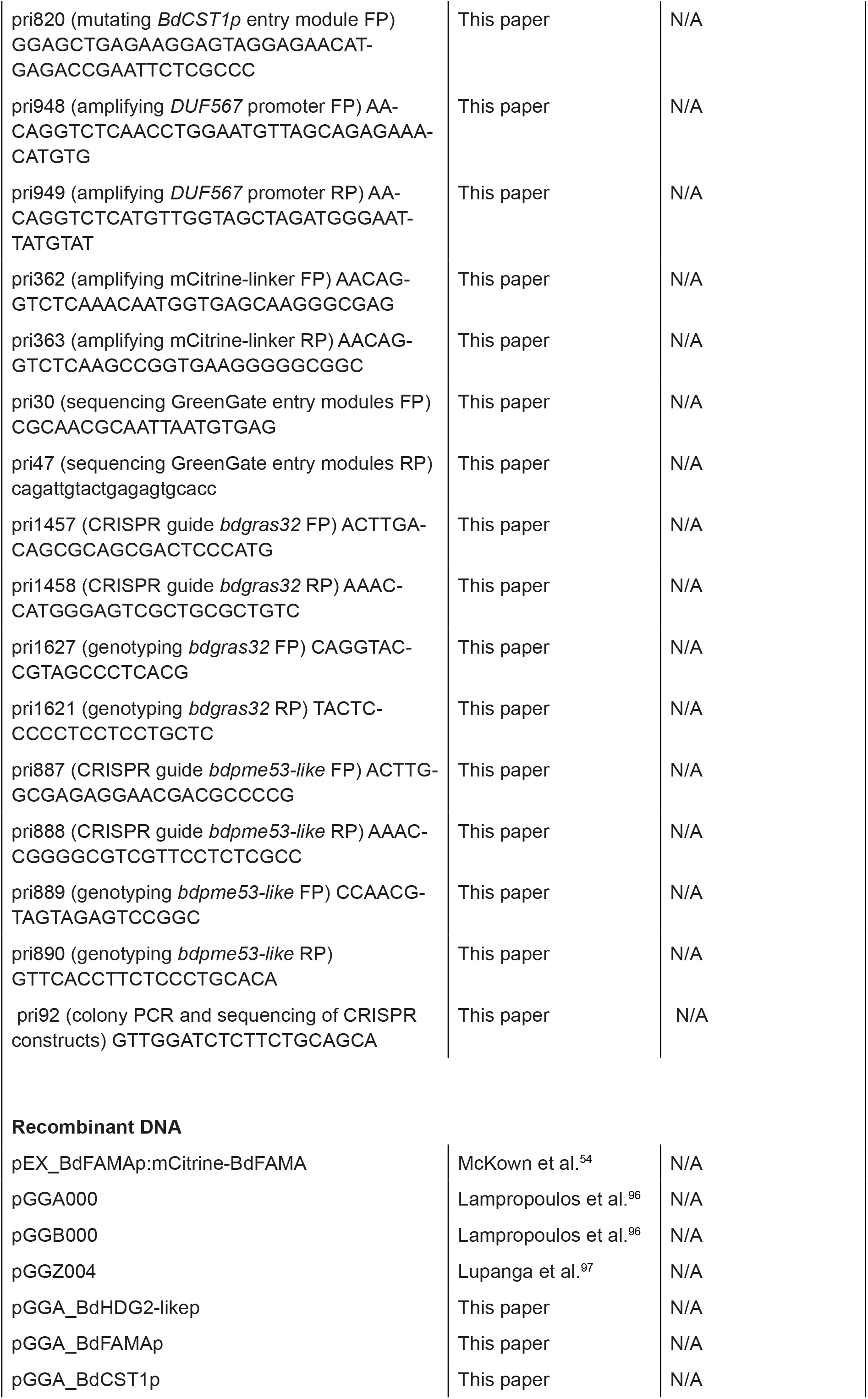
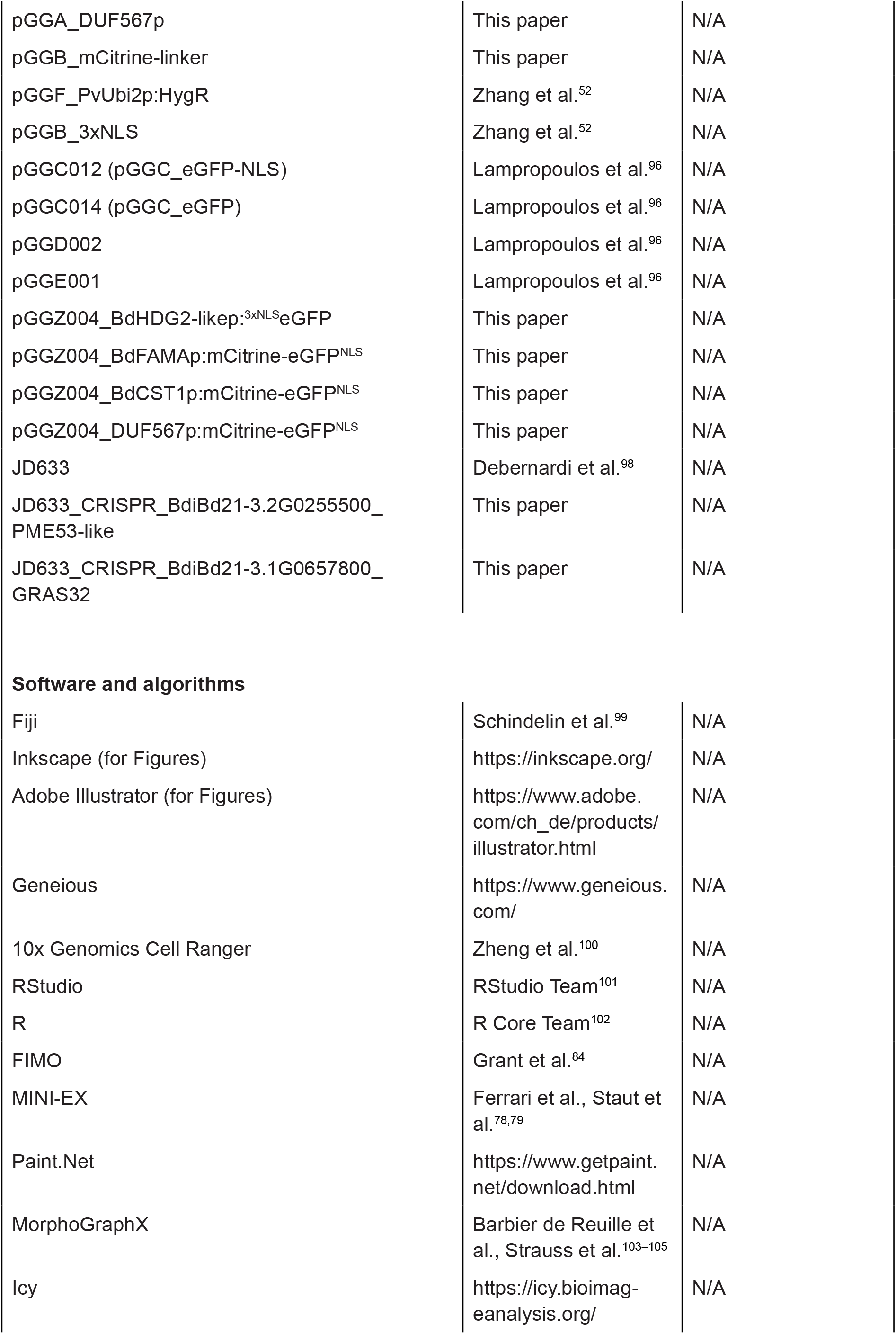

## Resource availability

### Lead contact

Further information and requests for resources and reagents should be directed to and will be fulfilled by the lead contact Michael T. Raissig (michael.raissig@unibe. ch).

## Materials availability

Recombinant DNA, transgenic reporter lines and mutant lines are available upon request.

## Data and code availability

Single-cell RNA-sequencing and bulk RNA-sequencing data have been deposited at GEO and are publicly available (GEO: GSE307277 and GSE306756). The expression browser of the single-cell RNA-sequencing dataset can be found online at https://shiny.ips.unibe.ch. All quantitative data generated and analyzed in this study can be found in the supplementary materials. Microscopy images reported in this paper will be shared by the lead contact upon request. R scripts used to analyze the datasets in this study, as well as a step by step protocol for protoplasting and HCR RNA-FISH of *B. distachyon* leaf tissue can be found on Github: https://github.com/raissig-lindner-lab/Berg-et-al_2025_ScSeq. Any additional information required to reanalyze the data reported in this paper is available from the lead contact upon request.

## Experimental model details

### Plant material and growth conditions

The *Brachypodium distachyon* Bd21-3 ecotype was used as wild type (WT) and the *sid* (also known as *bdmute-1*) line first described in ^9^ was used as the mutant line when creating the single-cell transcriptomic atlas. Seeds were dehusked and then vernalized and stratified for 2 days in water (dark, 4°C). Then, they were planted on soil (four parts Einheitserde CL ED73 (Einheitserdewerke, Werkverband e.V., Sinntal-Altengronau, Germany) and one part Vermiculite) and grown in a greenhouse or growth chamber with a 18 h light:6 h dark cycle (photosynthetically active photon flux density (PPFD) 200-400 μmol m^−2^ s^−1^; day temperature 28°C, night temperature 22°C) as described in ^106^.

For the imaging of *B. distachyon* transcriptional reporter lines *DUF567p:mCitrine-eGFP*^*NLS*^ and *BdFAMAp:m-Citrine-eGFP*^*NLS*^, as well as the *bdpme53-like* mutant plants for phenotyping, were grown on soil as described above. *BdHDG2-likep:*^*3XNLS*^*eGFP, BdCST1p:mCitrine-eGFP*^*NLS*^ and the *bdgras32* mutant plants (plus their wild type control) were grown on MS plates (Murashige & Skoog (Duchefa Biochemie B.V, Haarlem, The Netherlands), 1% Agar (w/v), pH 5.7; ^107^) and imaged 1 week (reporter) or 6 days (mutant) after germination.

*Hordeum vulgare* (barley) cultivar Golden Promise plants for the smRNA-FISH experiment were grown as described in ^27^. To reiterate: Plants were grown in soil (Einheitserde CL ED73) in 96-well trays with 4g/L Osmocote Exact Hi.End 3-4M, 4th generation (ICL Group, Ladenburg, Germany) under long day conditions (16 h light at 20°C and 8 h dark at 16°C). Grains were pre-germinated at 4°C for 4 days on a layer of wet paper in Petri dishes and vegetative shoot apical meristems were collected 11 days after sowing in soil.

For the barley reporter line, *HvFCP1p:VE-NUS-H2B*^56^, which is in the barley cultivar Golden Promise Fast^108^ background, plants were grown under the same conditions as described above, with the pre-germination treatment lasting 3 days.

## Method details

### Generation of protoplasts and single-cell RNA-sequencing

Single cells (protoplasts) were extracted from two tissue types of 3 week old, soil-grown plants. 7 samples were generated from the leaf developmental zone for 5 WT and 2 *sid* samples collected at different dates and locations (details on the samples in Table S1, Table S2). For this, 15 young, not yet unrolled leaves were carefully pulled from the enveloping sheath (Fig. 1B), each leaf from a different plant individual, and the lowest 3-5 mm were cut and collected in a petri dish with 3 ml Milli-Q water on ice. 2 WT samples were used to generate the vegetative shoot apex with leaf primordia of 8 and 10 individuals per sample, respectively. The plant was cut off right above the soil level and the older leaves were carefully removed under a binocular microscope to reach the shoot apical area. A cut was made right below the youngest outgrowing leaf as indicated with a dashed line in Fig. 1C and the tissue piece was collected in a petri dish with 3 ml milli-Q water on ice. Protoplasts were generated using a modified version of the protocol described in ^109^, with detailed steps as follows: The samples were vacuum infiltrated in a vacuum desiccator for 5 min and then incubated for 45 min to hydrate the cell walls (RT, dark, 90 rpm shaking). In the meantime, enzymes (Cellulase “Onozuka RS” (Duchefa, final concentration 2.4% (w/v)) and Macerozyme R-10 (Duchefa, final concentration 0.4% (w/v))) were added to 3 ml of the washing solution (D-Mannitol (Roth, final concentration 0.4 M), MES hydrate (Sigma-Aldrich, St. Louis, Missouri, USA, final concentration 20 mM), KCl (Sigma-Aldrich, final concentration 20 mM), CaCl_2_ (Sigma-Aldrich, final concentration 10 mM), BSA (Sigma-Aldrich, final concentration 0.1% (w/v), pH adjusted to 5.7 with 1 M Tris-HCl (pH8)). After incubation, the water was removed and replaced by 3 ml enzyme solution and then vacuum-infiltrated for 5 min. Then the sample was cut into very small pieces using nail scissors (less than 1 mm) and incubated for 90 min (RT, dark, 90 rpm shaking). About 3 mm of the tips of 1 ml pipette tips were cut off and the protoplast suspension was gently pipetted up and down for 3 min to mechanically release protoplasts. Then the protoplast-containing solution was transferred (cut pipette tip!) and slowly filtered through a 20 µm mesh filter (CellTrics, Sysmex cat. no. 04-004-2325) into a 1.5 ml LoBind tube (Eppendorf, Basel, Switzerland) (on ice). The sample was centrifuged for 3 min (300xg, 4°C). Two washing steps followed where the supernatant was removed with an uncut pipette tip without disturbing the pellet, before careful resuspension in 1 ml washing solution and centrifuged again for 3 min (300xg, 4°C). After two rounds of washing, the pellet was resuspended in 500 ml washing solution for the developmental zone samples and 100 ml for the vegetative shoot apex samples. Protoplast viability was assessed by staining with fluorescein diacetate (FDA (Honeywell International Inc., Fluka™, 0.01% (w/v) solution in acetone) for 30 min in the dark (only done for cells that were not used for sequencing). Green fluorescence of viable cells was visualized using an epifluorescence microscope (Leica DM5000B or DM2000 (Leica Microsystems, Wetzlar, Germany)). Cells were counted using a Fuchs-Rosenthal chamber (Brand GmbH + Co KG, Wertheim, Germany) on a DM2000 or DM5000b light microscope (Leica Microsystems) and brought to the sequencing facility for quality control, library preparation and sequencing (Deep Sequencing Core Facility, Heidelberg University, Germany and Next Generation Sequencing Platform, University of Bern, Switzerland respectively). The Chromium Next GEM Single Cell 3’ Reagent Kit v3.1 (10x Genomics, Pleasanton, USA) was used for library preparation and the libraries were sequenced using different Illumina Short Read sequencing platforms (see Table S1).

### Data preparation and primary analysis of the single-cell RNA-sequencing data

The Illumina sequencing files were first pre-processed using Cell Ranger (10X Genomics Cell Ranger v7.0.1, ^100^) with default settings. In short, the reference genome *Brachypodium distachyon* Bd21-3 v1.2 (obtained from Phytozome: https://phytozome-next.jgi.doe.gov/info/BdistachyonBd21_3_v1_2) was prepared with Cell Ranger *mkref* and afterwards the libraries were de-multiplexed, barcodes were processed and gene counts were computed with Cell Ranger *count*. For the 2022 datasets, both libraries (WT and *sid*) were sequenced twice using different sequencing platforms (see Table S1) and the libraries were merged using Cell Ranger *count*. Then, the data was filtered to exclude cells with high ambient RNA content using the SoupX package with default settings (version 1.6.2, ^110^) in RStudio and the SoupX objects were used to initialize Seurat objects for downstream processing with the Seurat package (version 5.1.0, ^111–115^). The following steps were repeated for each of the 9 datasets: To filter low quality cells, those with more than 5% mitochondrial and/or more than 10% chloroplast counts were excluded (mitochondrial and chloroplast genes used in this context are based on blast results from *Arabidopsis thaliana* genes (^116^; *B. distachyon* gene accessions list on GitHub: https://github.com/raissig-lindner-lab/Berg-et-al_2025_ScSeq), as well as cells with less than 1250 or more than 50’000 UMIs (nCount_RNA) and less than 500 or more than 10’000 features (nFeature_RNA). Within-cell expression was normalized with NormalizeData (default settings). FindVariableFeatures (default settings) was used to identify the top 2000 variable genes in the dataset. Gene expression was normalized across cells with ScaleData (default settings). A principal component analysis (PCA, RunPCA for 50 principal components) followed by Uniform Manifold Approximation and Projection (UMAP, RunUMAP with 30 dimensions) was conducted to reduce dimensionality of the datasets. Potential doublet cells were filtered out using the DoubletFinder package (version 2.0.4, ^117^) according to the GitHub instructions (https://github.com/chris-mcginnis-ucsf/DoubletFinder). Expected doublet formation rates were estimated using the Chromium Next GEM Single Cell 3’ Reagent Kit v3.1 User Guide as reference (10x Genomics, https://assets.ctfassets.net/an68im79xiti/1eX2FPdpeCgnCJtw4fj9Hx/7cb84edaa9eca04b607f9193162994de/CG000204_ChromiumNextGEMSingleCell3_v3.1_Rev_D.pdf). All 9 datasets were subsequently merged without integration to preserve the distinct tissue identity of shoot apices and developing leaves. The resulting final dataset was again processed using FindVariableFeatures, ScaleData and RunPCA (100 principal components). FindNeighbors (40 dimensions), FindClusters (resolution 0.8) and RunUMAP (40 dimensions) then yielded 29 clusters. G2-to-M (G2/M) and S phase genes were identified by blasting *A. thaliana* cell cycle genes (^118^; *B. distachyon* gene accessions list on GitHub: https://github.com/raissig-lindner-lab/Berg-et-al_2025_ScSeq) and cell cycle states were assigned using the CellCycleScoring function. Tissue identities were assigned to clusters by comparing expression patterns of known marker genes (see Fig.1, Fig. S1, Fig. S2, Table S3). To obtain the epidermal and stomatal cell file subsets, clusters were extracted and re-clustered according to the above described pipeline. To optimize the epidermal subset, an iterative process of manual removal of subclusters that do not express epidermal marker genes and re-clustering of the remaining cells was employed (further details in the R Script available on GitHub). For the stomatal lineage subset, epidermal clusters were annotated using stomatal lineage marker genes and extracted from the epidermal dataset. Cell types/stages were identified and annotated using known marker genes (see Fig. 2-5). Visualization of the single-cell sequencing data was done using Seurat and ggplot2 (version 3.5.2, ^119^). The complete R script is available on GitHub: https://github.com/raissig-lindner-lab/Berg-et-al_2025_ScSeq.

### Pseudotime analysis

For the pseudotime analysis of the guard cell lineage, the clusters 2, 5-7, 9 and 12 of the stomatal subset were extracted and re-clustered (Fig. S4B, S5A,B). For the hair cell (HC) lineage, clusters 6, 7, 10-13, 15 and 19 were extracted from the epidermal subset and re-clustered as well (Fig. S3B, S5D,E). The resulting Seurat objects were converted to monocle 3 cell_data_set objects using the SeuratWrappers package (version 0.4.0, ^120^). Monocle 3 (version 1.3.7, ^121–123^) was used to process the datasets (dimension = 50, resolution = 0.001 for the GC lineage, 0.0001 for the HC lineage), to identify and visualize pseudotime trajectories and compute and visualize the expression of selected marker genes along the pseudotime trajectories. The R script is available on GitHub: https://github.com/raissig-lindner-lab/Berg-et-al_2025_ScSeq.

### Gene regulatory network and GO term analysis

Gene regulatory network analysis of the stomatal lineage subset was conducted using MINI-EX (version 3.1, ^78,79^). First, input files needed to be created for *Brachypodium distachyon* Bd21-3 genome v1.2. The expression matrix, cluster markers and annotations were extracted from the Seurat object of the stomatal lineage subset as described on the MINI-EX github page (https://github.com/VIB-PSB/MINI-EX/blob/main/docs/data_preparation.md). The list of *B. distachyon* transcription factors (TFs) was downloaded from the Plant Transcription Factor Data-base: https://planttfdb.gao-lab.org/index.php?sp=Bdi. The .meme file and .txt information file for *B. distachyon* TF motifs were also downloaded from the Plant Transcription Factor Database: https://planttfdb.gao-lab.org/download. php#bind_motif. A conserved bHLH TF motif (CANNTG, ^85,86^) for the stomatal bHLH TFs BdSPCH1, BdSPCH2, Bd-MUTE, BdFAMA, BdICE1 and BdSCRM2 was manually added to the list. To map the TF motifs to the genome, the *B. distachy* Bd21-3 genome v1.2 .fasta file was down-loaded from Phytozome (https://phytozome-next.jgi.doe.gov/info/BdistachyonBd21_3_v1_2). The FIMO online tool (https://meme-suite.org/meme/tools/fimo, ^84^) was used to look for TF binding sites (TFBS) across the genome which were then manually filtered to obtain a list of target genes (TGs). FIMO TFBS hits across each chromosome were linked to genes by using the chromosomal position information which is available in the FIMO results and can be linked to the gene position given in the *B. distachyon* Bd21-3 v1.2 .gff file. Only one hit per TG and TF was kept and the thus cleaned TG list was extracted and used as input for MINI-EX.

After all required input files were obtained, MINI-EX was run with default settings and including the motif filtering analysis (the motif filter argument was set to “TF-F-motifs” to use the motifs known for a TF family). Further analysis and visualization of the MINI-EX output files was done in RStudio. The top 5 TFs per cluster were filtered based on their cluster-specific Borda ranking in the MINI-EX rankedRegulons output file. To compare the targetomes of BdMUTE and BdFAMA, the MINI-EX edgeTable output file was filtered for TGs in cluster 7 and then for those that could be bound by BdMUTE or BdFAMA. The TF-exclusive genes were selected by filtering for genes only bound by either of the TFs. By calculating the ratio of weight_BdMUTE_ (as an indicator of the strength of co-expression between the TF and the TG) by weight_BdFAMA_ and considered TGs with a weight ratio > 3 as preferentially regulated by Bd-MUTE and a weight ratio < −3 indicated those genes that may rather be regulated by BdFAMA. The list of TGs that were classified as preferentially regulated by BdMUTE was merged with the BdMUTE-exclusive TGs and the same was done for BdFAMA. Gene Ontology (GO) terms for biological processes obtained using the R packages Go.db (version 3.20.0, ^124^) and AnnotationDbi (version 1.68.0, ^125^) were linked to the TGs and for the top 25 GO terms per dataset, their frequency was visualized with ggplot2. Any additional information and details on file preparation and modifications, as well as analysis and visualization of the MINI-EX output files can be found in the GRN R Script on Github (https://github.com/raissig-lindner-lab/Berg-et-al_2025_ScSeq).

### Selection of candidate genes

Differential gene expression analysis was conducted in R to find marker genes of different cell types and/ or developmental stages using the Seurat FindMarkers function (default settings). Thus obtained candidates were screened manually by checking their gene expression feature plots. Of this selection, only genes were considered for downstream analysis that were not upregulated (fold change > 3) in the protoplasted bulk RNAseq data compared to non-protoplasted (see below). A subset of those genes representing a diverse panel of cell types and developmental stages were chosen for HCR RNA-FISH and transcriptional reporters to confirm their expression and thus cluster annotation *in planta*. The selected genes were linked to known orthologues and functionally annotated using information obtained from Phytozome gene annotations^126^, InterPro^127^, PaperBLAST^128^ and the OMA browser^129^.

### Bulk RNA-sequencing datasets to compare protoplasted and non-protoplasted tissue

To assess gene expression changes induced by protoplasting, total RNA was extracted from both protoplasted and non-protoplasted developmental leaf zones of 3 weeks old Bd21-3 (WT) plants. Protoplasts were prepared from 10 developmental zones according to the protocol above and 5 leaf zones were used for the non-treated (i.e. non-protoplasted) control (3 biological replicates per condition). RNA extraction was performed with the RNeasy micro kit (QIAGEN GmbH, Hilden, Germany) with on-column DNAse digest following the manufacturer’s instructions. RNA concentration was assessed with the Qubit RNA High Sensitivity (HS) Assay Kit (Invitrogen, Thermo Fisher Scientific, Massachusetts, USA). RNA quality check, library preparation and sequencing on a NextSeq 550 platform (Illumina Inc., San Diego, USA) was conducted at the Deep Sequencing Core Facility of the Heidelberg University (Germany).

Pre-processing of the bulk RNA-seq results was done on https://usegalaxy.org/. After quality control of the sequencing results with FastQC^130^, the bulk transcriptomes were mapped to the Bd21-3 v1.1 genome (https://phytozome-next.jgi.doe.gov/info/BdistachyonBd21_3_v1_1) with bowtie2^131–133^. Further processing was done using R (version 4.5.1, ^102^) in RStudio (version 2024.12.01, ^101^). Read counts were calculated with summarizeOverlap (version 1.42.0, ^134^) and subsequently analyzed with DeSeq2 (version 1.46.0, ^135^) to compare protoplasted and non-protoplasted samples. Finally, gene expression was normalized by transcripts per million (TPM). A volcano plot showing differentially expressed genes (DEGs) was generated with ggplot2. Genes above a log2 fold change of 3 were considered upregulated and those below −3 downregulated (Fig. S3C).

### HCR RNA-FISH

Leaf developmental zones of young, not yet unrolled leaves of 3 weeks old soil-grown plants were harvested and hairpin chain reaction RNA fluorescence in situ hybridization (HCR™ RNA-FISH (v3.0), Molecular Instruments, Inc., Los Angeles, USA; ^62,63^) was conducted as described in ^53^. Two fluorophores were used, one with signal in the green range (B1 amplifier with fluorophore 488) and one with signal in the red range (B2 amplifier with fluorophore 594). For each gene, between 3 and 20 probes were designed by Molecular Instruments and one color fluorophore was assigned (color in the images depicted in this study corresponds to the respective fluorophore). For ease of reference, the protocol is included again here: The protocol used here was adapted from ^136,137^ and spans 5 days from material collection to final confocal imaging. On day 1, young, not yet unrolled, leaves were collected and the bottom 5-8 mm was harvested in a Petri dish in a few drops of a fixative solution (Fixative FAA; formaldehyde solution [final concentration 4% (v/v)] (Sigma-Aldrich), glacial acetic acid [final concentration 5% (v/v)] (Thermo Fisher Scientific), absolute ethanol [final concentration 50% (v/v)] in nuclease-free water (Thermo Fisher Scientific)). The leaf piece was cut once in longitudinal and once in transverse direction and then collected in a 2 ml tube with Fixative FAA and this was repeated for four to five additional leaves that were added to the same sample tube. After sample collection, vacuum was applied to the tube several times until all leaf pieces were submerged and then incubated at room temperature (RT) for 3 h. Fixative FAA was replaced by a series of ethanol concentrations (10%, 30%, 50%, 70%) and, for each concentration, the samples were microwaved five times for 30 s at 180 W^137^. After the last microwaving step with 70% ethanol, the sample was stored in a −20°C freezer (the sample can be stored like this for several weeks).

On day 2, the sample was allowed to warm up to RT and then rehydrated through a series of washes at RT on a tube revolver (Thermo Fisher Scientific): first with 50% ethanol/50% DPBS-T (Tween20 [final concentration 0.1% (v/v)] (Sigma-Aldrich) in Dulbecco’s phosphate-buffered saline (DPBS) (Gibco, Thermo Fisher Scientific, Massachusetts, USA)) for 15 min and then twice with 100% DPBS-T for 15 min each step. Aliquots of Proteinase K solution [1 M Tris-HCl, pH 8 (final concentration 0.1 M), 0.5 M EDTA, pH 8 (final concentration 0.05 M), Proteinase K (final concentration 80 μg/ml) (Thermo Fisher Scientific) in nuclease-free water] had been prepared and frozen at −20°C and one aliquot was taken out to thaw at RT during the first incubation step with 100% DPBS-T. The DPBS-T was replaced by the Proteinase K solution and incubated by applying vacuum for 5 min and then digesting for 25 min at 37°C on the thermomixer (Eppendorf). During this step, the sample was agitated every 5 to 10 min. Subsequently, the sample was washed twice for 15 min in DPBS-T at RT on the tube rotator (Eppendorf). The second fixative solution (Fixative II; formaldehyde solution [final concentration 4% (v/v)] in DPBS-T) replaced the DPBS-T and the sample was again vacuum-infiltrated for 10 min followed by 20 min at RT on the tube rotator. In the meantime, two 500 μl aliquots of the HCR™ Probe Hybridization Buffer (Molecular Instruments) were prepared: one was left to reach RT and the other was put to 37°C in the thermomixer. The sample was washed twice for 15 min in DPBS-T at RT on the tube rotator and then incubated in the RT aliquot of the Probe Hybridization Buffer by applying vacuum for 10 min and then prehybridized for 1 h at 37°C in the thermomixer with shaking (1000 rpm). The probe solution was prepared by adding 0.8 pmol (i.e. 2 μl of the 1 μM stock) of each HCR™ probe to 500 μl of the Probe Hybridization Buffer at 37°C. For multiplexing of probes, two probes that can be linked to different amplifiers were added to the same tube. As much as possible was removed from the pre-hybridization solution while being careful to not pick up the leaf pieces and the final probe solution was added to the sample before an overnight (22 h) incubation step at 37°C in the thermomixer with shaking (1000 rpm).

To prepare for day 3, an aliquot of HCR™ Probe Wash Buffer (Molecular Instruments) was warmed up to 37°C. Additionally, two aliquots of HCR™ Amplification Buffer were prepared and warmed up to RT: one with 250 μl and one with 500 μl. The excess probes were washed out by washing four times with 500 μl Probe Wash Buffer for 15 min at 37°C in the thermomixer with shaking (1000 rpm). Another two washing steps with SSC-T buffer (20x SSC buffer [final concentration 25% (v/v)] (Invitrogen, Thermo Fisher Scientific), Tween20 [final concentration 0.1% (v/v)] in nuclease-free water) for 5 min each at RT in the thermomixer with shaking (1000 rpm) were carried out and the rest of the SSC-T buffer was stored at 4°C. Next, the hairpins h1 and h2 for the amplifier B1 with fluorophore 488 and amplifier B2 with fluorophore 594 (Molecular Instruments) were put in an ice bucket to slowly thaw. In the meantime, the SSC-T buffer was replaced by 500 μl Amplification Buffer and vacuum was applied for 10 min before 50 min pre-amplification at RT on the tube rotator. During this incubation step, 6 pmol of hairpins h1 and h2 (i.e. 5 μl of the 3 μM stocks) were put in separate tubes (2 tubes for each amplifier), heated at 95°C for 90 s and then kept in a dark drawer at RT for 30 min. Then all hairpins were assembled in the prepared tube of 250 μl Amplification Buffer. As much as possible of the pre-amplification solution was removed while trying to avoid damage to the leaf pieces and then the Amplification Buffer with hairpins h1 and h2 was added to the sample and incubated for about 42 h in the dark at RT.

On day 5, the SSC-T buffer was taken from the 4°C fridge to warm up to RT. Excess hairpins were removed from the sample by washing with SSC-T buffer in a series of washes: twice for 5 min, twice for 30 min and once for 5 min. To visualize cell walls, the sample was stained with SCRI Renaissance Stain 2200 (SR2200, Tokyo Future Style, Inc., Tsukuba City, Japan, [final concentration 0.001% (v/v)] in SSC-T buffer) or Calcofluor White (Sigma-Aldrich, [final concentration 0.0001% (w/v)] in SSC-T buffer) for 1 min and then washed three times with SSC-T buffer. For the imaging, a Leica Stellaris 5 (Leica Microsystems) was used with the following settings: SR2200 was excited at 405 nm excitation, green probe signal (fluorophore 488) was excited at 490 nm, red probe signal (fluorophore 594) was excited at 586 nm and all emissions were captured within the range of the respective fluorophore emission peaks.

Subsequently, color channels were merged and adjusted for brightness and contrast in Fiji^99^.

### Cloning of *Brachypodium distachyon* transcriptional reporter constructs

Transcriptional reporter constructs were generated using the Greengate cloning system^96^. All promoter sequences were amplified from genomic DNA (CTAB extraction method, ^138^) of wild type (WT) *B. distachyon* Bd21-3 using the Q5 polymerase (New England Biolabs (NEB), Ipswich, USA). Promoter sequences were selected upstream of the coding sequences annotated in the *B. distachyon* Bd21-3 v1.2 genome (https://phytozome-next.jgi.doe.gov/info/BdistachyonBd21_3_v1_2). The length of promoters depended on the distance to the next gene upstream of the gene of interest and was also adjusted slightly to optimize primers. The *BdHDG2-like* promoter (*BdHDG2-likep*) was 3’584 bp and amplified using the primer pair pri121/pri122. For *BdFAMA*, the promoter (*BdFAMAp*) was 4’472 bp long and amplified with primers pri454/pri475. The *BdCST1* promoter (*BdCST1p*) was amplified in two fragments using the primers pri117/pri289 (fragment 1) and pri288/pri118 (fragment 2) to remove a BsaI site in the promoter. After ligating the two amplicons, the final amplicon was lacking a piece of the UTR, which was added by amplifying the entry module with pri820/819 and using insertional site-directed mutagenesis with the missing sequence attached to the forward primer (pri820) to complete the entry module, leading to a final promoter length of 3’555 bp. For *DUF567*, the promoter (*DUF567p*) was 1’480 bp and amplified using primers pri948/pri949. For the B module containing mCitrine with a linker, the mCitrine-linker sequence was amplified from *BdFAMAp:mCitrine-BdFAMA* ^54^ using the primers pri362/363. Amplicons were PCR-purified using the NucleoSpin Gel and PCR Clean-up kit (Macherey-Nagel AG, Oensingen, Schweiz) and digested with BsaI (NEB) or its isoschizomer Eco31I (Thermo Fisher Scientific). The pGGA000 (backbone for promoters) and pGGB000 (backbone for mCitrine-linker) entry vectors were digested with BsaI/Eco31I and dephosphorylated using Antarctic Phosphatase (NEB). Then, the respective inserts were ligated into the respective entry vector using the T4 ligase (Thermo Fisher Scientific) overnight at 16°C creating pGGA_BdHDG2-likep, pGGA_BdFAMAp, pGGA_BdCST1p, pGGA_DUF567 and pGGB_mCitrine-linker plasmids. Plasmids were transformed into *E. coli* and after overnight incubation, a colony PCR with pri30/pri47 was conducted to confirm insert lengths. Positive colonies were grown overnight to amplify the plasmid before extraction using the NucleoSpin Plasmid kit (Macherey-Nagel AG) and final sequencing to confirm the correct plasmid.

The *PvUbi2p*-driven hygromycin resistance cassette was amplified from pTRANS_250d^139^ and used to create a Greengate F module as described in ^52^.

Further Greengate entry modules pGGB_3xNLS (3xNLS), pGGC012 (eGFP^NLS^), pGGC014 (eGFP) were generously provided by Prof. Dr. Karin Schumacher’s group and pGGD002 (D-dummy) and pGGE001 (rbcS terminator) were provided by Prof. Dr. Jan Lohmann’s group^96^.

The final transcriptional reporter constructs were created using the GreenGate protocol as described in ^96^. For ease of reference, the protocol is included again here: The GreenGate entry modules (pGGA_specific-promoter; pGGB_3xNLS or pGGB_mCitrine-linker; pGGC_eGFP or pGGC_eGFP-NLS; pGGD_dummy; pGGE_rbcS-terminator and pGGF_hygromycin-resistance) were repeatedly digested and ligated into the pGGZ004^97^ for 50 cycles (each cycle 5 min at 37°C for digestion and 5 min at 16°C for ligation), followed by enzyme inactivation steps at 50°C (5 min) and 80°C (5 min). The final ligations were transformed into *E. coli* for selection and amplification and verified by enzymatic digestion profiles. Correct assembly of the various inserts was further confirmed by sequencing (whole-plasmid sequencing or Sanger sequencing of the overhangs).

### Design and cloning of CRISPR constructs

CRISPR guides were selected using the “Find CRISPR sites” tool of Geneious Prime (version 2024.0.3, GraphPad Software LLC d.b.a Geneious, activity scoring based on ^140^). The selected guides and their reverse complement with added cloning overhangs for *BdGRAS32* (pri1457 and pri1458) and *BdPME53-like* (pri887 and pri888) were annealed and phosphorylated using the T4 PNK (NEB). The JD633 plasmid^98^ was used as CRISPR backbone and first digested using the PaqCI enzyme (NEB), followed by de-phosphorylation using Antarctic Phosphatase (NEB). The annealed guides and the backbone were ligated overnight at 16°C using the T4 DNA ligase (NEB). Competent *E. coli* NEB 5-alpha (NEB) were transformed with the CRISPR plasmids and amplified overnight. On the next day, colony PCR with a primer binding on the CRISPR backbone (pri92) and the respective reverse guide was conducted to identify positive colonies, which were then grown overnight to amplify the plasmid. Plasmids were isolated with the NucleoSpin Plasmid kit (Macherey-Nagel AG) and sent for sequencing using pri92 to check for correct insertion of the guide.

### *Brachypodium distachyon* plant transformation

To prepare the constructs for transformation, *Agrobacterium tumefaciens* AGL1^95^ were transformed with the respective plasmids and grown on plates overnight at 28°C. Positive colonies were used for transformation of embryonic calli of wild type *B. distachyon* Bd21-3 following the plant transformation protocol described in ^52^. For ease of reference, the protocol is included again here: Young, transparent embryos were isolated and grown for three weeks at 28°C in the dark on callus induction media (CIM; per L: 4.43 g Linsmaier & Skoog basal media (LS; Duchefa), 30 g sucrose, 600 ml CuSO_4_ (1 mg/ml, Sigma/Merck), 500 ml 2,4-D (5 mg/ml in 1M KOH, Sigma/Merck), pH 5.8, plus 2.1 g of Phytagel (Sigma/Merck)). After incubation, crisp, yellow callus pieces were subcultured to fresh CIM plates and further incubated at 28°C in the dark for two weeks. Then, calli were broken down to 2-5 mm small pieces, transferred to fresh CIM plates and subcultured for one more week at 28°C in the dark. For transformation, agrobacteria with the desired construct were dissolved in liquid CIM media (same media as above without the phytagel) with freshly added 2,4-D (2.5 mg/ml final concentration), Acetosyringone (200 mM final concentration, Sigma/Merck), and Synperonic PE/F68 (0.1% final concentration, Sigma/Merck). The OD600 of the agrobacteria solution was adjusted to 0.6. Around 100 calli were incubated for at least 10 min in the bacteria solution, dried off on sterile filter paper and incubated for three days at room temperature in the dark. After incubation, transformed calli were moved to selection media (CIM + Hygromycin (40 mg/ml final concentration, Roche, Basel, Switzerland) + Timentin (200 mg/ml final concentration, Ticarcillin 2Na & Clavulanate Potassium (Duchefa))) and incubated for one week at 28°C in the dark. Then, calli were moved to fresh selection plates and incubated for two more weeks at 28°C in the dark. Next, calli were moved to callus regeneration media (CRM; per L: 4.43 g of LS, 30 g maltose (Sigma/Merck #M5885), 600 ml CuSO4 (1 mg/ml), pH 5.8, plus 2.1 g of Phytagel). After autoclaving, cool down and add Timentin (200 mg/ml final concentration), Hygromycin (40 mg/ml final concentration), and sterile Kinetin solution (0.2 mg/ml final concentration, Sigma/Merck). Calli were incubated at 28°C with a 16h light:8h dark cycle (70-80 mmol PAR m^−2^ s^−1^). After 1-6 weeks in the light, shoots will form. Shoots that are longer than 1 cm and ideally have two or more leaves were moved to glass cups for rooting (Weck glass and packaging GmbH, Bonn, Germany) containing rooting media (per L: 4.3 g Murashige & Skoog including vitamins (Duchefa), 30 g sucrose, adjust pH to 5.8, add 2.1 g Phytagel. After autoclaving, cool down and add Timentin (200 mg/ml final concentration)). Once roots had formed, plantlets were moved to soil (consisting of 4 parts Einheitserde CL ED73 (Einheitserdewerke, Werkverband e.V., Sinntal-Altengronau, Germany) and 1 part Vermiculite) and grown in a greenhouse or growth chamber with 18h light:6h dark cycle (250-350 mmol PAR m^−2^ s^−1^). For the first 2-3 days, trays with the transgenic plantlets would still be covered with a transparent lid to maintain humidity.

### Imaging of abaxial mature epidermis

To obtain the image of the abaxial leaf epidermis in Fig. 2A, lignin of a mature leaf was stained with Basic Fuchsin (Sigma-Aldrich) and imaged together with cell wall autofluorescence as described in ^49^. For ease of reference, the protocol is included again here: Small leaf fragments previously fixed and cleared in 7:1 ethanol:acetic acid were washed in 70% ethanol and then transferred to distilled water with 0.02% (v/v) Tween20 (Sigma-Aldrich) for rehydration for 3 hours. Samples were incubated in 30 ml of 0.01% Basic Fuchsin for 5 min and washed twice with 30 ml of 50% (v/v) glycerol for 5 min (2.5 min per washing step) and then mounted in 50% (v/v) glycerol for imaging. Samples were imaged on a Stellaris 5 confocal microscope (Leica Microsystems). Cell wall autofluorescence was excited at 405 nm and emissions captured in the range of 490-550 nm. Basic Fuchsin was excited at 561 nm and emissions were captured between 573-603 nm. Stacks of 0.33 µm steps were obtained. For the image in Fig. 2A, a sum of slices Z-projection was performed in Fiji^99^.

### Imaging of reporter lines

For *DUF567p:mCitrine-eGFP*^*NLS*^ and *BdFAMAp:m-Citrine-eGFP*^*NLS*^, the developmental zone of the youngest, not yet unrolled leaf of 2-3 weeks old plants was imaged. For *BdHDG2-likep:*^*3xNLS*^*eGFP* and *BdCST1p:mCitrine-eGFP*^*NLS*^, second leaves of plate-grown seedlings were imaged 5-7 days after germination. Cell walls were stained with propidium iodide (PI (Invitrogen, Thermo Fisher Scientific), 0.5% (v/v)) for 3-5 min. A Leica Stellaris 5 confocal microscope (Leica Microsystems) with the following settings was used to visualize the different channels: PI was excited at 549 or 584 nm, mCitrine was excited at 515 nm and eGFP was excited at 489 nm. Emissions were captured in the range of the emission peak of the respective fluorophore. Subsequently, color channels were merged and adjusted for brightness and contrast in Fiji^99^. For the *BdCST1* reporter, z-stacks were taken and Maximum Projection images were created.

Imaging of the barley reporter line *HvFCP1p:VE-NUS-H2B*^56^ was performed with the confocal microscope Zeiss LSM880 (Carl Zeiss AG, Oberkochen, Germany), with Plan-Apochromat 20×/0.8 or Plan-Apochromat 40x/1 objectives. Fresh barley shoot apical meristems and leaf primordia were stuck on a double-sided adhesive tape on an objective slide, stained with PI (0.3 mM) for 3 min, washed three times with water and subsequently covered with a cover slide before being placed under the microscope. PI was excited at 561 nm and Venus at 514 nm. Signal was collected within the fluorophore’s emission peak. Pictures were analysed using Fiji^99^.

### *Hordeum vulgare* multiplex smRNA-FISH

smRNA-FISH (Molecular Cartography) of the *Hordeum vulgare* (barley) cultivar Golden Promise was conducted as described in the associated publication^27^. For ease of reference, the protocol is included again here: Vegetative shoot apical meristems and surrounding leaf primordia were fixed overnight in phosphate buffered saline (PBS) containing 4% paraformaldehyde (PFA) and 0.03% Triton X-100 (Sigma-Aldrich), dehydrated through an ethanol series, and embedded under vacuum in Paraplast Extra (Leica Biosystems, Deer Park, USA). 10 μm sections of the embedded tissues were placed on glass slides (Resolve Biosciences), deparaffinized, gradually rehydrated, and permeabilized with proteinase K (Thermo Fisher Scientific). After re-fixation, acetylation and dehydration through an ethanol series, the slides were mounted with SlowFade-Gold Antifade Mountant (Invitrogen, Thermo Fisher Scientific). Probe hybridisation and imaging were performed by Resolve Biosciences. Image analysis was performed in Fiji^99^ using the Polylux plugin (Polylux_V1.9.0) from Resolve Biosciences.

### Genotyping of CRISPR mutant lines

To genotype the CRISPR mutant lines, genomic DNA was extracted from T0 and T1 transgenic plants using the CTAB extraction method^141^. Genotyping primers were selected up- and downstream of the CRISPR guide site to amplify the putative mutation site (*bdgras32*: pri1627 and pri1621, *bdpme53-like*: pri889 and pri890). The mutation site was PCR-amplified and the correct size of the amplicon was confirmed in a standard 1% agarose gel. After PCR purification with the NucleoSpin Gel and PCR Cleanup kit (Macherey-Nagel AG), the PCR product was sent for sequencing with the added primer pri1621 (for *bdgras32*) or pri889 (for *bdpme53-like*) to identify or confirm the mutation and the zygosity.

### Phenotyping of *bdgras32* mutant line

6 days after germination, second leaves of plate-grown seedlings of wild type (WT) and *bdgras32* (T1 generation) were imaged on a Stellaris 5 confocal microscope (Leica Microsystems). Cell walls were stained using propidium iodide (PI, 0.5% (v/v)) for 4 minutes. PI was excited at 549 nm and emissions were captured around the emission peak. Images were taken of cell files around developmental stage 6 to show the misdivision phenotype, and around the first asymmetric epidermal divisions (i.e., developmental stage 1 to 2) for subsequent analysis. Stitching of images was done in Fiji^99^ using the pairwise stitching tool^142^ and Paint.NET (version 5.1.9, https://www.getpaint.net/index.html). Then, stitched sequences were cropped to frames of the same length (388.93 µm) with the first asymmetric divisions approximately in the center (see Fig. 7I). 2D segmentation of 5 cell files including one stomatal lineage cell file was conducted in MorphoGraphX (version 2.0.1-379, ^103–105^) as follows: Images were adjusted with Brighten/Darken (value = 3) and the signal was projected onto the mesh (minimum distance 0, maximum distance 2) before auto segmentation. Mistakes were corrected by manual tracing and watershed segmentation until a correct cell mesh was created. Only cells were kept that are fully visible in the frame. Corner triangles were fixed, the mesh was smoothened (walls only, 3 passes) and then Heatmap Classic (manual range 0-300) was used to calculate and visualize cell areas (Fig. 7I). Statistical significance analysis using unpaired, two-sided Student’s *t* test (stats package version 4.3.1, ^102^) and visualization of results was done in R. The R script for mutant analysis as well as the data table output of MorphoGraphX is available on Github: https://github.com/raissig-lindner-lab/Berg-et-al_2025_ScSeq.

### Phenotyping of *bdpme53-like* mutant line

Leaf-level gas exchange measurements of wild type (WT) and *bdpme53-like* (T1 generation) plants were conducted and subsequently analyzed as described in ^143^. For ease of reference, the protocol is included again here: Physiological parameters for each genotype were obtained from 3-4 week old soil-grown, non-flowering plants. The youngest, fully expanded leaf was measured using the LI-6800 Portable Photosynthesis System with the 6800-01A Multiphase Flash Fluorometer chamber (LI-COR Biosciences Inc., Lincoln, NE, USA) in the 2 cm^2^ chamber. Environmental conditions were programmed as described in ^106^: flow rate, 500 µmol s^−1^; fan speed, 10000 rpm and leaf temperature, 28°C. Gas exchange in reaction to changing light intensities was measured at relative humidity, 40% and CO_2_ concentration, 400 µmol mol^−1^. Values were automatically logged every 60 seconds with 20 minute steps for the light intensities in units of photosynthetically active radiation (PAR), 1000 (high light) −100 (low light) – 1000 −0 (darkness) PAR m^−2^ s^−1^. PAR is equivalent to Photosynthetic photon flux density (PPFD), which is in μmol m^−2^ s^−1^.

As *B. distachyon* leaves are not broad enough to fill the 2 cm^2^ chamber, data from each run was corrected manually in the Excel sheet by individual leaf area, which was calculated as leaf width multiplied by the chamber diameter. Subsequent averaging of values across all individuals per species, outlier removal, correction by stomatal density and final visualization was conducted in R Studio using the licornetics R package (version 2.1.2, ^143,144^) that is available on Github with further documentation (https://github.com/lbmountain/licornetics).

Leaves were collected after gas exchange measurements and fixed in 7:1 ethanol:acetic acid for at least 24h. Samples were rinsed in water and mounted on slides in Hoyer’s solution (70% (w/v) chloralhydrate (Roth, Karlsruhe, Germany), 4% (w/v) glycerol (Thermo Fisher Scientific), 5% (w/v) gum arabic (Roth)). The abaxial leaf side was imaged using a Leica DM2000 DIC microscope (Leica Microsystems). For stomatal density, 3 fields of view (image size 0.198 mm^2^) per leaf were counted in Fiji^99^, resulting in 29-53 stomata per individual and divided by the image size to obtain stomatal density per mm^2^. Statistical significance analysis using unpaired, two-sided Student’s *t* test (stats package version 4.3.1, ^102^) and visualization of results was done in R. The R script for mutant analysis is available on Github: https://github.com/raissig-lindner-lab/Berg-et-al_2025_ScSeq.

### *Brachypodium distachyon* developing leaf single-cell atlas web tool

To create an expression browser, we generated a template onto which gene expression can be plotted. First, we created the template file following the tutorial provided in the ggPlantmap Github: https://github.com/leonardojo/ggPlantmap/blob/main/guides/TutorialforXMLfile.pdf. In short, the Icy software (version 2.5.2.0, https://icy.bioimageanalysis.org/) was used to trace and create a template with outlines and labels for different tissues and cell types and save them as .xml. In R Studio, the ggPlantmap package (version 1.1.0, ^88^) was used to convert the .xml file to a ggPlantmap object. The Seurat AverageExpression function was used to extract mean expression of genes per cluster identity. For the main tissue assignments, the whole dataset was used and Seurat clusters were labelled for analysis as follows: Shoot apex = cluster 2; vasculature = clusters 3, 16, 19, 20, 21, 22, 26; mesophyll = clusters 0, 1, 5, 6, 8, 9, 14, 18, 24, 28; epidermis = clusters 4, 7, 11, 13, 17, 23, 25, 27. For annotation of hair cells (HCs) and silica cells, the epidermal subset was used and Seurat clusters were labelled for analysis as follows: Silica cells = clusters 0, 1, 18; early HCs = clusters 7, 10; middle HCs = clusters 6, 12; late HCs = cluster 13. For stomatal lineage cell types and stages, the stomatal lineage subset was used and Seurat clusters were labelled for analysis as follows: Early GMCs = cluster 9; dividing GMCs = clusters 7, 12; early GCs = cluster 6; late GCs = cluster 5; SCs = clusters 4, 13; interstomatal cells = clusters 1, 8. The ggPlantmap.heatmap function was then used to plot gene expression onto the previously created template.

The UMAP feature plots depicting gene expression were created as follows: The Seurat FetchData function was used to obtain UMAP locations of cells, their assigned identity and the expression of genes (features) in a given cell. Visualization of expression of a gene of interest was done using the ggplot2 package. The shiny package (version 1.9.1, ^145^) was used to create the website, which was then deployed with Docker (Docker Inc., Palo Alto, USA). Further packages required for the creation of the web tool: tidyverse (version 2.0.0, ^146^), Cairo (version 1.6-2, ^147^), bslib (version 0.8.0, ^148^), shinythemes (version 1.2.0, ^149^), shinycssloaders (version 1.1.0, ^150^), MetBrewer (version 0.2.0, ^151^) and arrow (version 20.0.0.2, ^152^).

The R script used to create the web tool (also works locally) is available on Github: https://github.com/raissig-lindner-lab/Berg-et-al_2025_ScSeq.

