## Supplementary material for "The developing leaf of the wild grass *Brachypodium distachyon* at single-cell resolution": Figs S1-S6; Tables S1-S3

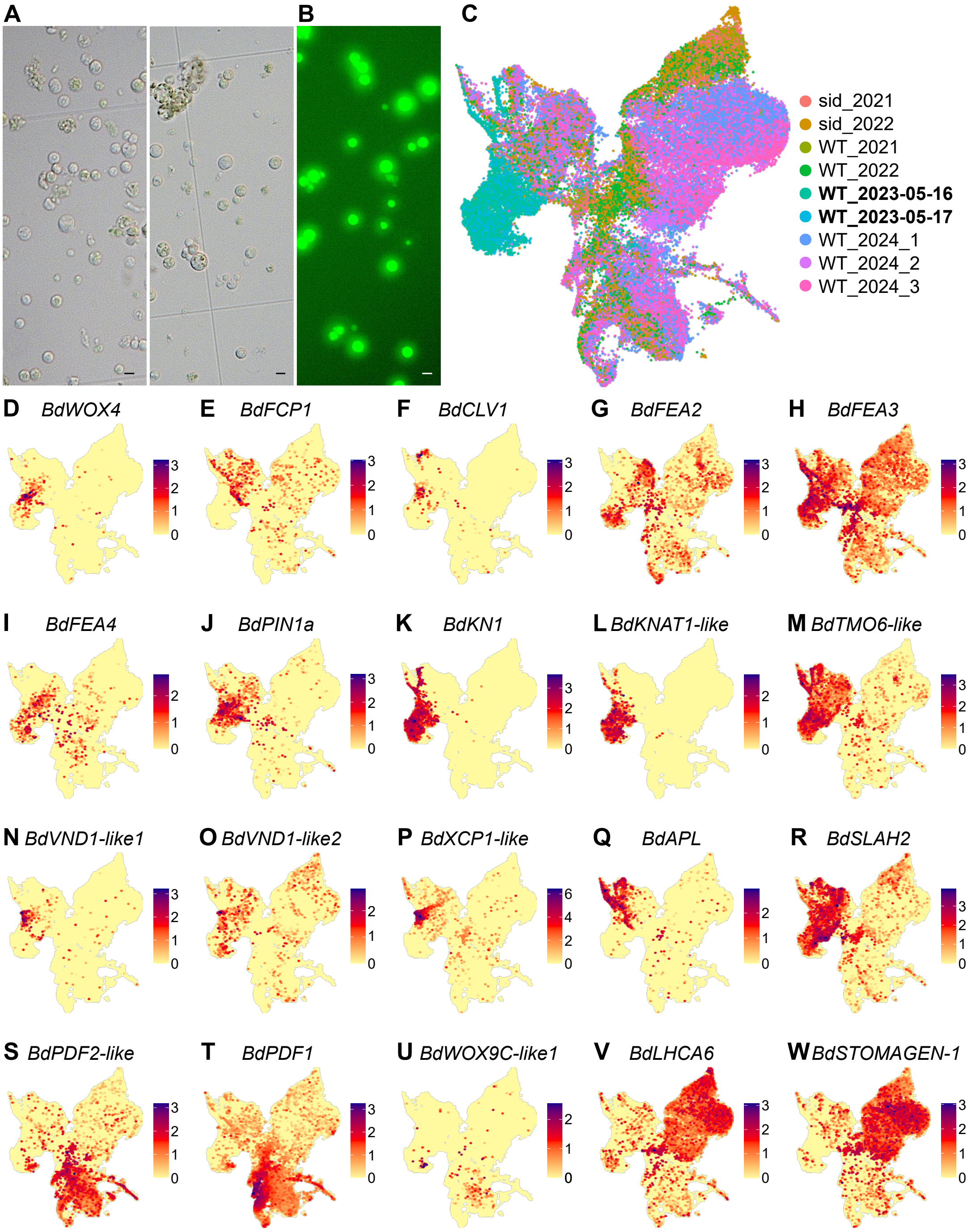


**Figure S1.** Related to Fig. 1. **(A)** DIC images of protoplasted cells. Scale bars, 10 µm. **(B)** Epifluorescence microscopy image of protoplasts stained with fluorescein diacetate (FDA). Viable cells glow green. Scale bar, 10 µm. **(C)** Whole dataset UMAP plot with color indicating the library origin. vSAM & primordia libraries in bold. **(D-R)** Whole dataset UMAP feature plots of marker genes for vegetative shoot apical meristem (vSAM) and early vasculature. *BdKNOTTED1* (K) and *BdSLOWLY ACTIVATING ANION CHANNEL (SLAC1) HOMOLOGUE 2* (*BdSLAH2*, R) are also shown in Fig. 1E with a different color scheme. **(S-U)** Whole dataset UMAP feature plots of marker genes for the leaf epidermis. *BdPROTODERMAL PATTERNING FACTOR 2-like* (*BdPDF2-like*; S) is also shown in Fig. 1E with a different color scheme. **(V, W)** Whole dataset UMAP feature plots of marker genes for the mesophyll. *BdSTOMAGEN-1* is also shown in Fig. 1E with a different color scheme. Each dot in the UMAP plots represents the transcriptome of a single cell. Color legends in the UMAP feature plots indicate expression strength.

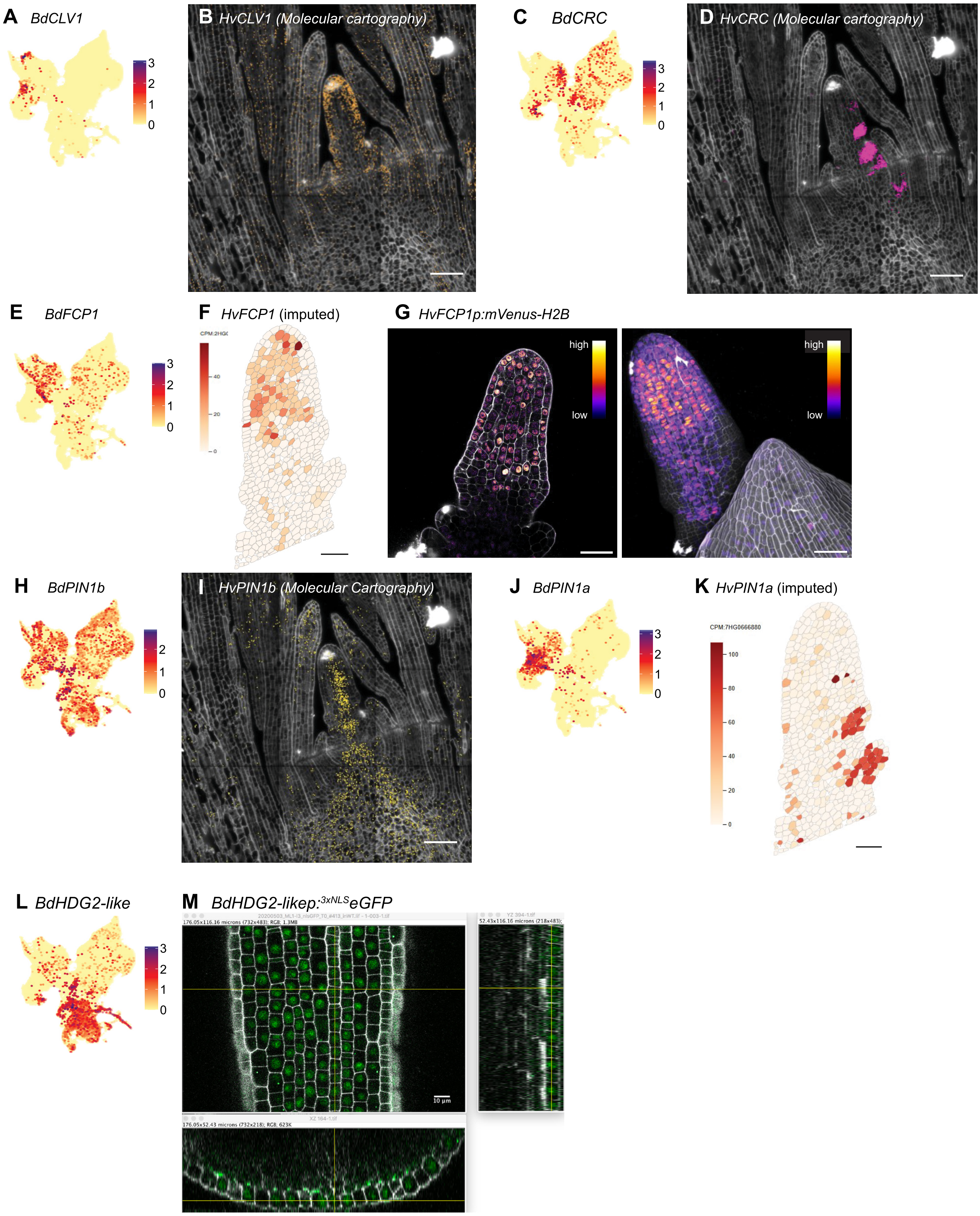


**Figure S2.** Related to Fig. 1. **(A)** Whole dataset UMAP feature plot of *BdCLAVATA1* (*BdCLV1*). Same plot as in Fig. S1F. **(B)** smRNA-FISH (Molecular cartography) image of barley *HvCLV1* in the shoot apex with early leaves. Full size version of the image shown in Fig. 1c of [56] with different color scheme. Also shown as a zoom-in without gene expression channel in Fig. 1B of [27]. **(C)** Whole dataset UMAP feature plot of *BdCRABS CLAW* (*BdCRC*). **(D)** smRNA-FISH image of barley *HvCRC* in the shoot apex with early leaves. Full size version of the image shown in Fig. 3F of [27] with different color scheme and only *HvCRC* expression. Also shown as a zoom-in without gene expression channel in Fig. 1b of [27]. **(E)** Whole dataset UMAP feature plot of *BdFON2-LIKE CLE PROTEIN 1* (*BdFCP1*). Same plot as in Fig. S1E. **(F)** Imputed expression of barley *HvFCP1*. Dataset and imputation method are described in [27]. **(G)** Expression of *HvFCP1p:Venus-H2B* in barley vegetative shoot apical meristem (vSAM, left) and leaf primordial epidermis (right, at Waddington stage 1-1.5). Color legend indicates expression strength. Cell walls stained with propidium iodide (PI, gray). **(H)** Whole dataset UMAP feature plot of *BdPINFORMED1b* (*BdPIN1b*). **(I)** smRNA-FISH image of barley *HvPIN1b* (also known as *HvPIN1*). Also shown as a zoom-in without gene expression channel in Fig. 1b of [27]. **(J)** Whole dataset UMAP feature plot of *BdPIN1a*. Same plot as in Fig. S1J. **(K)** Imputed expression of barley *HvPIN1a*. Dataset and imputation method are described in [27].

**Figure S2 (continuted). (L)** Whole dataset UMAP feature plot of *BdHOMEODOMAIN GLABROUS2-like* (*BdHDG2-like*). **(M)** Expression of *BdHDG2-likep:$^{3xNLS}$eGFP* in *B. distachyon* leaf epidermis (z-stack) with orthogonal sections indicated by yellow lines; transverse section below, longitudinal section to the right. Color in the smRNA-FISH images indicates expression of the gene of interest. Each dot in the UMAP plots represents the transcriptome of a single cell. Color legends in the UMAP feature plots and imputed data indicate expression strength. Scale bars, 50 µm, unless otherwise indicated.

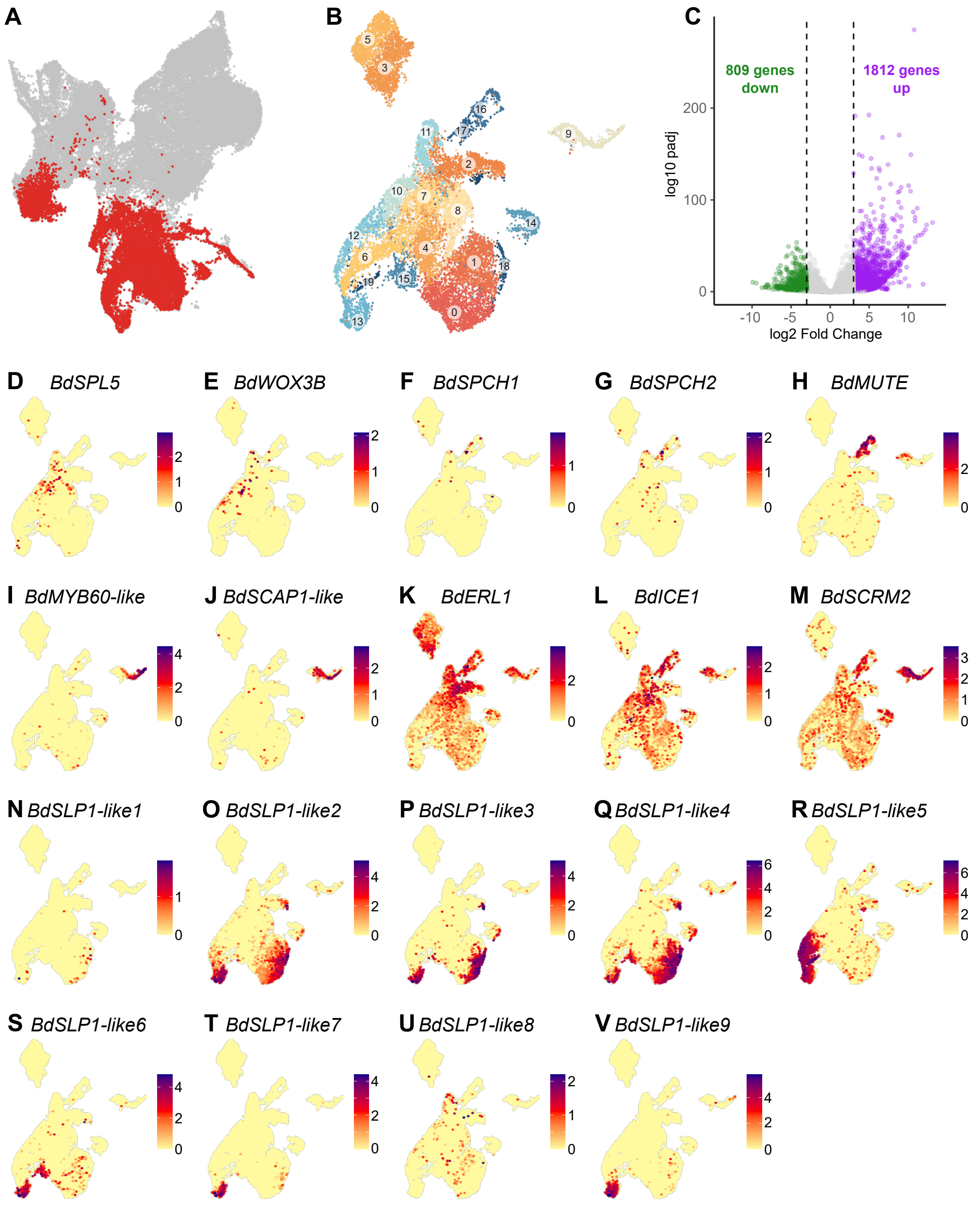


**Figure S3.** Related to Fig. 2. **(A)** Whole dataset UMAP plot with cells belonging to the epidermal subset highlighted in red. **(B)** UMAP plot of the epidermal subset with numbered Seurat clusters. **(C)** Volcano plot showing logarithmic fold change of genes in response to the protoplasting protocol. Genes are considered downregulated (green) if the log2fold change is < -3 and upregulated (magenta) if the fold change is > 3. **(D, E)** Epidermis UMAP feature plots of hair cell lineage marker genes. **(F-M)** Epidermis UMAP feature plots of stomatal lineage marker genes. **(N-V)** Epidermis UMAP feature plots of the *BdSILIPLANT1-like* (*BdSLP1-like*) family. *BdSLP1-like3* (P) and *BdSLP1-like5* (R) are also shown in Fig. 2L,M. *BdSLP1-like1* (N) is protoplasting-affected. Each dot in the UMAP plots represents the transcriptome of a single cell. Color legends in the UMAP feature plots indicate expression strength.

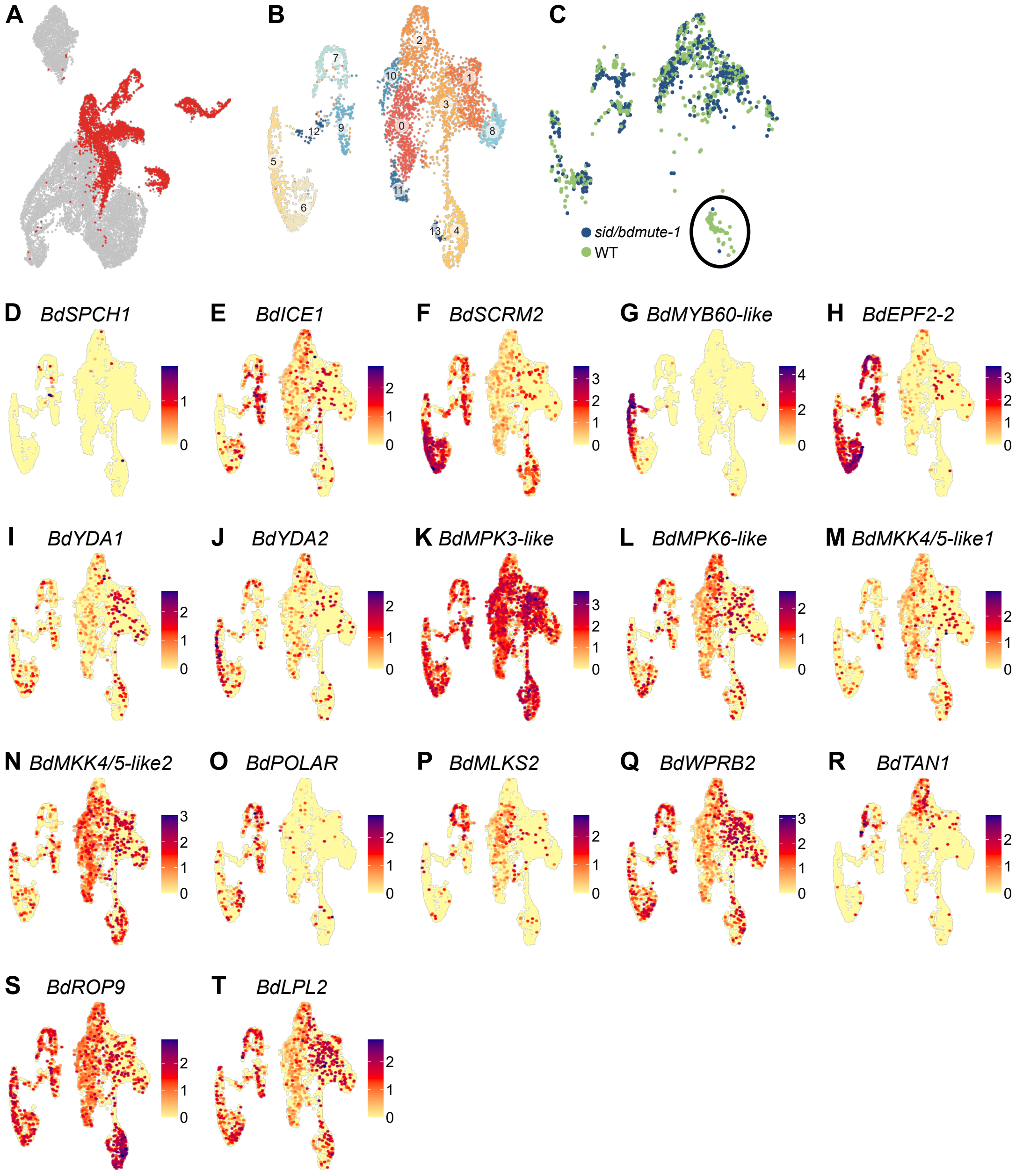


**Figure S4.** Related to Fig. 3 and 5. **(A)** Epidermis UMAP plot with cells belonging to the stomatal lineage subset highlighted in red. **(B)** UMAP plot of the stomatal lineage subset with numbered Seurat clusters. **(C)** Stomatal lineage UMAP plot with color indicating genotype. Same UMAP plot as in (B), but only the two wild-type and two *sid/bdmute-1* libraries generated at the same time are shown (i.e., the two 2021 and the two 2022 datasets, Fig. S1C). The subsidiary cell (SC) clusters enriched for wild type (WT) cells are indicated by a circle. **(D-H)** Stomatal lineage UMAP feature plots of stomatal lineage marker genes. **(I-N)** Stomatal lineage UMAP feature plots of *MITOGEN-ACTIVATED PROTEIN* (*MAP*) kinase genes potentially involved in stomatal development. **(O-T)** Stomatal lineage UMAP feature plots of genes involved in the subsidiary mother cell (SMC) division of grasses. Each dot in the UMAP plots represents the transcriptome of a single cell. Color legends in the UMAP feature plots indicate expression strength.

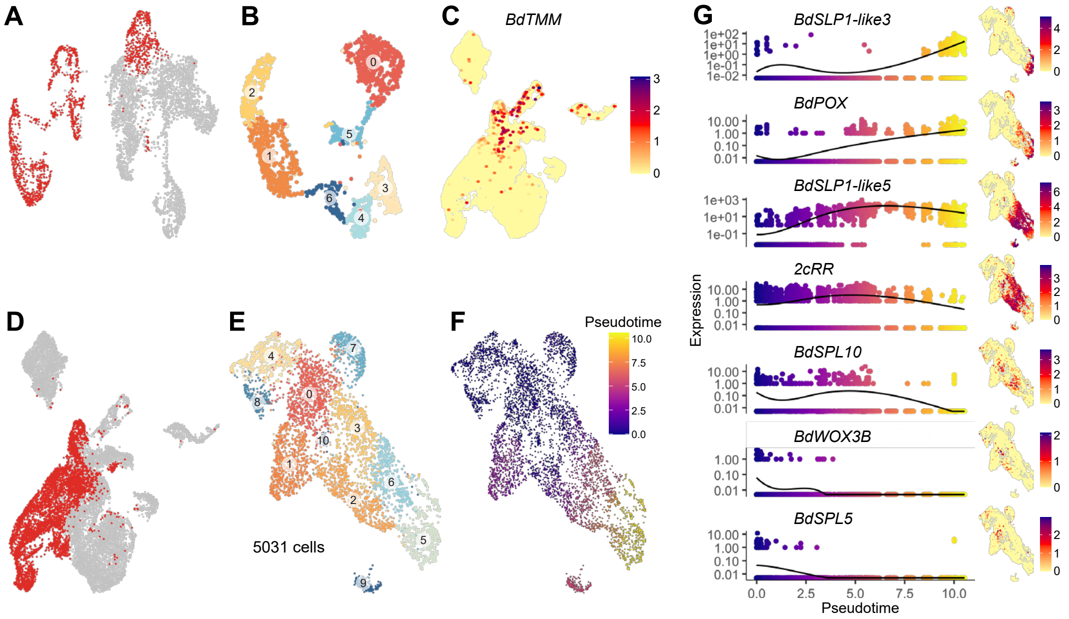


**Figure S5.** Related to Fig. 4. **(A)** Stomatal lineage UMAP plot of the stomatal lineage subset with cells belonging to the guard cell (GC) lineage subset highlighted in red. **(B)** UMAP plot of the GC lineage subset with numbered Seurat clusters. **(C)** Epidermis UMAP feature plot of *BdTOO MANY MOUTHS* (*BdTMM*). **(D)** UMAP plot of the epidermal subset with cells belonging to the hair cell (HC) lineage highlighted in red. **(E)** HC lineage UMAP plot with numbered Seurat clusters. N = 5’032 cells. **(F)** UMAP plot of the HC lineage subset with color indicating pseudotime. **(G)** Dot plot showing expression of marker genes across the HC lineage subset pseudotime gradient. UMAP feature plots of the respective genes are shown on the right. Color legends in the UMAP feature plots indicate expression strength.

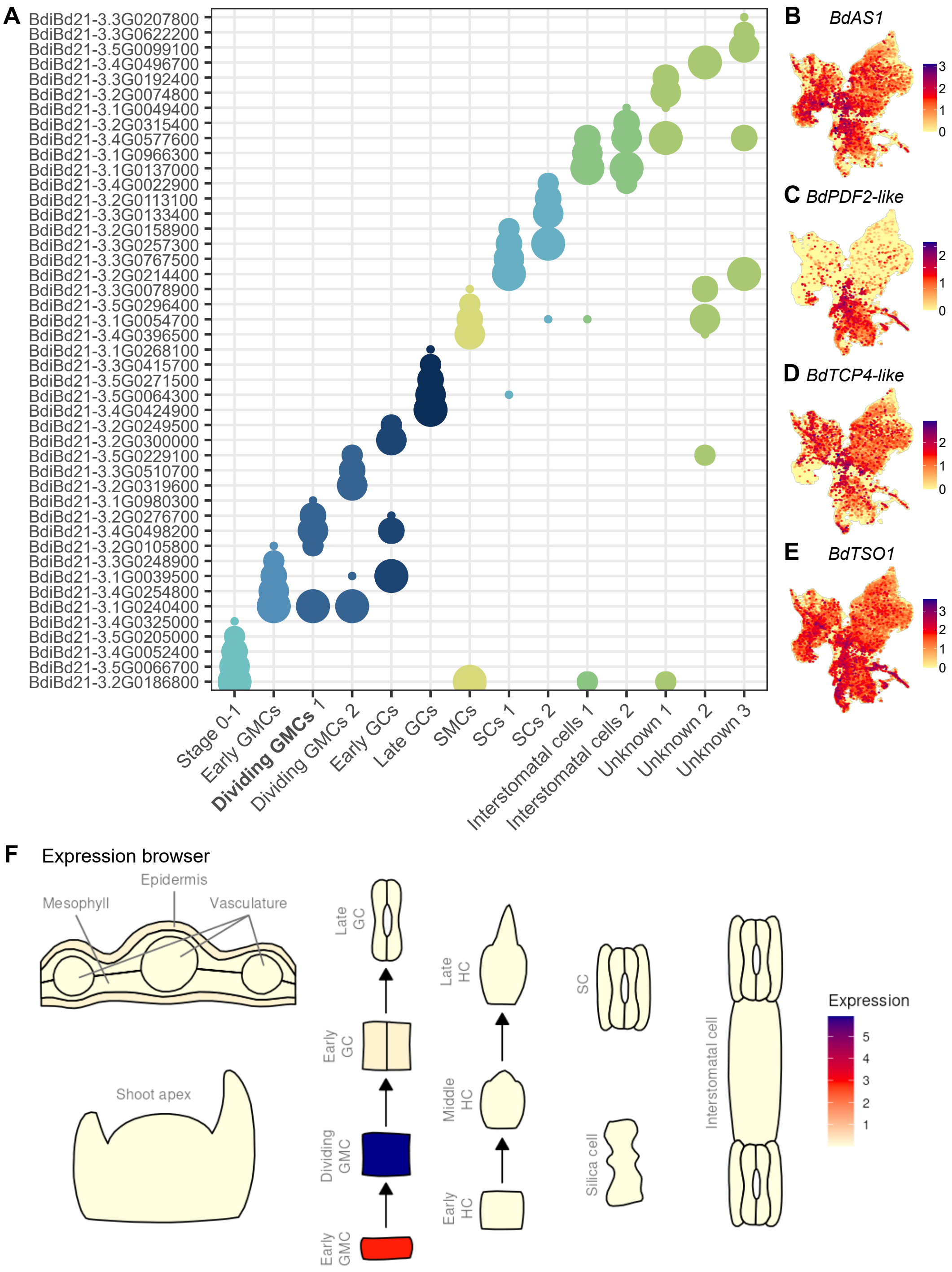


**Figure S6.** Related to Fig. 6. **(A)** Dot plot showing the top 5 transcription factors (TFs)/regulons per cluster. Same plot as in Fig. 6A, here with *B. distachyon* Bd21-3 gene accession numbers instead of names. **(B)** Whole dataset UMAP feature plot of *BdASYMMETRIC LEAVES 1* (*BdAS1*). **(C)** Whole dataset UMAP feature plot of *BdPROTODERMAL PATTERNING FACTOR 2-like* (*BdPDF2-like*). Same plot as in Fig. S1S and also shown in Fig. 1E, there with a different color scheme. **(D)** Whole dataset UMAP feature plot of *BdTEOSINTE BRANCHED 1, CYCLOIDEA, PROLIFERATING CELL FACTOR1/2 4* (*BdTCP4-like*). **(E)** Whole dataset UMAP feature plot of *BdTSO1*. **(F)** Representative image of the expression overview map which is accessible for any *B. distachyon* Bd21-3 gene on the website introduced in this study (here shown for *BdMUTE*). Each dot in the UMAP plots represents the transcriptome of a single cell. Color legends in the UMAP feature plots indicate expression strength.

**Table S1.** General overview of the scSeq datasets.

| **Dataset** | **Tissue** | **Location** | **Nr. of cells after filtering** | **Mean/median nr. of UMI** | **Mean/median nr. of features** | **GEO identifier** |
|---|---|---|---|---|---|---|
| WT 2021 | Leaf developmental zone | Heidelberg (DE) | 2’656 | 12’929/11’594 | 3’674/3’894 | GSM9220335 |
| sid 2021 | Leaf developmental zone | Heidelberg (DE) | 3’427 | 10’256/8’515 | 3’189/3’186 | GSM9220336 |
| WT 2022 | Leaf developmental zone | Heidelberg (DE) | 6’904 | 14’224/12’112 | 4’049/4’171 | GSM9220337 and GSM9220339 |
| sid 2022 | Leaf developmental zone | Heidelberg (DE) | 7’437 | 15’201/13’354 | 4’112/4’250 | GSM9220338 and GSM9220340 |
| WT 2023-05-16 | vSAM & leaf primordia | Heidelberg (DE) | 4’995 | 5’055/4’194 | 2’302/2’153 | GSM9220341 |
| WT 2023-05-17 | vSAM & leaf primordia | Heidelberg (DE) | 2’674 | 4’798/3’938 | 2’230/2’085 | GSM9220342 |
| WT 2024-1 | Leaf developmental zone | Bern (CH) | 13’441 | 5’087/3’956 | 2’223/2’039 | GSM9220343 |
| WT 2024-2 | Leaf developmental zone | Bern (CH) | 11’546 | 6’304/5’366 | 2’720/2’626 | GSM9220344 |
| WT 2024-3 | Leaf developmental zone | Bern (CH) | 16’607 | 6’734/5’955 | 2’841/2’763 | GSM9220345 |

**Table S2.** Necessary reported information to allow evaluation and repetition of a plant single cell/nucleus experiment, template from Grones et al. 2024

| | **Details** | **Experimental information** |
|---|---|---|
| **Biological material** | Species | *Brachypodium distachyon* |
| | Accession | Bd21-3 |
| | Genotype | WT and *sid/bdmute-1* |
| | Tissue type | Leaf developmental zones and vegetative shoot apex with leaf primordia |
| | Detailed growth conditions | Greenhouse or growth chamber, 18 h light: 6 h dark (day temperature 28°C, night temperature 22°C), PPFD 200-400 µmol $m^{-2}$ $s^{-1}$, soil-grown (four parts Einheitserde CL ED73, 1 part Vermiculite), 2 days vernalization (in water, 4°C, dark) |
| | Harvest conditions | 3-week-old plants, morning, 15 leaf developmental zones or 8-10 shoot apices |
| **Sample preparation** | Isolation protocol | Tissue was cut mechanically, incubated in water, digested using Cellulase “Onozuka RS” and Macerozyme R-10, and carefully pipetted up and down to release protoplasts |
| | Tissue dissection | Young, not yet unrolled leaves were pulled out from the enveloping older leaf and lowest 3-5 mm were cut off with a scalpel to harvest developmental zones. Shoot apices were dissected and harvested using a dissecting scope, forceps and a scalpel |
| | Fixation | Unfixed tissue |
| | Cell enrichment | Not applicable |
| | Total sample preparation time | 4 to 4.5 h (from harvest to submission for 1-2 samples at a time, time includes counting of cells at the microscope and time needed to dilute to final concentration),<br>2.5 to 3 h (from digestion start to submission for 1-2 samples at a time) |
| | Estimated cell number loaded | 12k cells (2021 datasets), 16.5k cells (2022 datasets), 18k cells (2023 and 2024 datasets) |
| | Instrument/Method/Kit | 10x Genomics Chromium Single Cell 3’ v3.1 |
| | Cell viability test | Fluorescein diacetate staining |
| **Libraries** | Library construction | GEM wells were prepared to encapsule protoplasts in droplets, cDNA was extracted per protoplast and libraries were prepared according to the guidelines provided by 10x Genomics (single cell 3' version 3.1) |

| | | |
|---|---|---|
| | End bias | 3' end |
| **Sequence results** | Instrument/method | NextSeq 550 (2021 datasets, first sequencing run of the 2022 datasets),<br>NextSeq 2000 (second sequencing run of the 2022 datasets, 2023 datasets),<br>Illumina NovaSeq 6000 (2024 datasets) |
| | Library layout/paired-end | Paired-end |
| | N° sequenced reads Reads/cell | ~32-54k (2021 datasets),<br>~75-84k (2022 datasets),<br>~93-154k (2023 datasets),<br>~24k-29k (2024 datasets) |
| **Raw data** | Reference genome | https://phytozome-next.jgi.doe.gov/info/BdistachyonBd21_3_v1_2 |
| | Annotation version | B. distachyon Bd21-3 v1.2 |
| | Mapping method (incl. software, customized settings) | 10x Genomics Cell Ranger v7.0.1 |
| | Mapping efficiency | > 91% to > 97% |
| | Sequencing saturation | 38-54% (2021 datasets),<br>42-45% (2022 datasets),<br>84-90% (2023 datasets),<br>54-61% (2024 datasets) |
| | Estimation of ambient RNA (Fraction reads in cell) | > 66% (2021 datasets),<br>> 71-77% (2022 datasets),<br>> 82-89% (2023 datasets),<br>> 75-85% (2024 datasets) |
| | Imputation method and settings | Not applicable |
| **Processed data** | N° captured cells (before filtering) | 5'480 (WT 2021),<br>6'023 (sid/bdmute-1 2021),<br>10'023 (WT 2022),<br>11'438 (sid/bdmute-1 2022),<br>7'600 (WT 2023-05-16),<br>4'285 (WT 2023-05-17),<br>22'531 (WT 2024-1),<br>18'565 (WT 2024-2),<br>22'828 (WT 2024-3) |
| | N° high quality cells | 69'686 |
| | Filter criteria: % mitochondrial reads/cell | < 5% |
| | Filter criteria: % chloroplast reads/cell | < 10% |

| | | |
|---|---|---|
| | Filter criteria: Minimum N° UMI/cell | > 1’250 and < 50’000 UMI;<br>> 500 and < 10’000 features |
| | N° total detected transcripts | > 91% (35’584 out of 39’068 genes) |
| | Doublet rate | 6% (2021 datasets), 8% (2022 datasets), 10% (2023 and 2024 datasets) |
| | Replicate comparisons | Bulk RNA-seq protoplasted vs. non-protoplasted tissues |
| | Batch correction method for merging (incl. reasoning for batch correction) | Not applicable |
| | Additional processing | High ambient RNA filtering (SoupX), doublet removal (DoubletFinder) |
| **Validation** | Method of automatic annotation of clusters | Not applicable |
| | Method of manual annotation (markers, gene function info) | Marker genes (*B. distachyon*, orthologs to genes known from other species; see Table S3) |
| | Verification in planta (e.g. Number of markers used for validation) | 13 marker genes verified with Hairpin Chain Reaction (HCR) RNA-fluorescence in situ hybridization,<br>4 marker genes verified with transcriptional reporter lines |
| **Data availability** | Analysis scripts & codes (GitHub) | Scripts available on Github: https://github.com/raissig-lindner-lab/Berg-et-al_2025_ScSeq |
| | Excel Tables DEG for each cluster | Supplementary information of the publication, Github: https://github.com/raissig-lindner-lab/Berg-et-al_2025_ScSeq |
| | Objects/count matrix in repository (which one, where?) | Single-cell data on GEO: GSE307277 |
| | On-line tool/browser URL | https://shiny.ips.unibe.ch/ |
| | Cell-level metadata table | Available in the R Script on Github: https://github.com/raissig-lindner-lab/Berg-et-al_2025_ScSeq |
| **Additional** | additional comments from the authors | Intronic reads were considered in Cell Ranger 7.0.1 |

**Table S3.** List of genes mentioned in the main text or figures of this paper.

| Name | Bd21-3 accession | Bradi accession | Related gene in other species | Relevant literature |
|---|---|---|---|---|
| *2cRR* | BdiBd21-3.3G0655900 | Bradi3g49440 | N/A | This paper |
| *BdAPL* | BdiBd21-3.3G0071200 | Bradi3g05500 | *AtAPL* | Bonke et al.[1] |
| *BdAS1* | BdiBd21-3.3G0192400 | Bradi4g03970 | *AtAS1, ZmRS2* | Tsiantis et al, Byrne et al.[2,3] |
| *BdCLV1* | BdiBd21-3.1G0402200 | Bradi1g30160 | *AtCLV1, HvCLV1* (HORVU.MOREX.r3.7HG0747230), *OsFON1, ZmTD1* | Clark et al., Suzaki et al., Bommert et al., Demesa-Arevalo et al., Vardanega et al.[4–8] |
| *BdCRC* | BdiBd21-3.1G0942300 | Bradi1g69900 | *HvCRC* (HORVU.MOREX.r3.4HG0396510) | Bowman and Smyth, Yamaguchi et al., Demesa-Arevalo et al.[7,9,10] |
| *BdCST1* | BdiBd21-3.2G0326500 | Bradi2g24850 | *ZmCST1* | Wang et al.[11] |
| *BdEPF2-1* | BdiBd21-3.5G0153600 | Bradi5g12220 | *AtEPF2, TaEPF2* | Hara et al., Hunt and Gray, Jangra et al.[12–14] |
| *BdEPF2-2* | BdiBd21-3.5G0306100 | Bradi5g23357 | *AtEPF2, TaEPF1* | Hara et al., Hunt and Gray, Jangra et al.[12–14] |
| *BdERECTA* | BdiBd21-3.1G0609800 | Bradi1g46450 | *AtERECTA* | Shpak et al., Herrmann and Torii, Chua and Lau[15–17] |
| *BdERL1* | BdiBd21-3.1G0662500 | Bradi1g49950 | *AtERL1* | Shpak et al., Herrmann and Torii, Chua and Lau[15–17] |
| *BdFAMA* | BdiBd21-3.2G0300000 | Bradi2g22810 | *AtFAMA, OsFAMA* | Ohashi-Ito and Bergmann, Liu et al., Wu et al., Wu et al., McKown et al.[18–21] |
| *BdFCP1* | BdiBd21-3.5G0168500 | Bradi5g13241 | *OsFCP1, HvFCP1* (HORVU.MOREX.r3.2HG0174890) | Ohmori et al., Vardanega et al.[4,22] |
| *BdFEA2* | BdiBd21-3.2G0427700 | Bradi2g34337 | *ZmFEA2, AtCLV2* | Je et al.[23] |
| *BdFEA3* | BdiBd21-3.2G0010000 | Bradi2g00920 | *ZmFEA3, HvFEA3* | Je et al., Demesa-Arevalo et al.[7,24] |
| *BdFEA4* | BdiBd21-3.1G0571300 | Bradi1g43900 | *ZmFEA4* | Pautler et al.[25] |
| *BdGELP1* | BdiBd21-3.2G0049500 | Bradi2g03807 | *ZmGELP1* | Sun et al.[26] |
| *BdGRAS32* | BdiBd21-3.1G0657800 | Bradi1g49630 | *OsDLT/GRAS-32* | Xie et al.[27] |
| *BdHDG2-like* | BdiBd21-3.3G0192600 | Bradi3g14500 | *AtHDG2* | Nakamura et al.[28] |
| *BdICE1* | BdiBd21-3.4G0254800 | Bradi4g17460 | *AtICE1, OsICE1, ZmICEb* | Kanaoka et al., Grimault et al. Raissig et al.[29–31] |
| *BdKN1* | BdiBd21-3.1G0135700 | Bradi1g10047 | *ZmKN1* | Smith et al.[32] |
| *BdKNAT1-like* | BdiBd21-3.1G0773000 | Bradi1g57607 | *AtKNAT1* | Lincoln et al.[33] |
| *BdLHCA6* | BdiBd21-3.4G0439300 | Bradi4g31257 | *AtLHCA6* | Jansson[34] |
| *BdMKK4/5-like1* | BdiBd21-3.1G0616400 | Bradi1g46880 | *AtMKK4/5* | Wang et al.[35] |
| *BdMKK4/5-like2* | BdiBd21-3.3G0709500 | Bradi3g53650 | *AtMKK4/5* | Wang et al.[35] |
| *BdMPK3-like* | BdiBd21-3.1G0885800 | Bradi1g65810 | *AtMPK3* | Wang et al.[35] |

| Name | Bd21-3 accession | Bradi accession | Related gene in other species | Relevant literature |
|---|---|---|---|---|
| *BdMPK6-like* | BdiBd21-3.1G0650300 | Bradi1g49100 | *AtMPK6* | Wang et al.[35] |
| *BdMUTE* | BdiBd21-3.1G0240400 | Bradi1g18400 | *AtMUTE, OsMUTE, ZmMUTE/BZU2* | Pillitteri and Torii, Raissig et al., Wang et al., Wu et al., Spiegelhalder et al.[21,36–39] |
| *BdMYB60-like* | BdiBd21-3.4G0234500 | Bradi4g16290 | *AtMYB60* | Cominelli et al.[40] |
| *BdPAN1* | BdiBd21-3.3G0526300 | Bradi3g39910 | *ZmPAN1* | Cartwright et al., Zhang et al.[41,42] |
| *BdPAN2* | BdiBd21-3.1G0783500 | Bradi1g58260 | *ZmPAN2* | Zhang et al.[43] |
| *BdPDF1* | BdiBd21-3.2G0082000 | Bradi2g06300 | *AtPDF1* | Abe et al.[44] |
| *BdPDF2-like* | BdiBd21-3.3G0204900 | Bradi3g15327 | *AtPDF2* | Abe et al.[45] |
| *BdPIN1a* | BdiBd21-3.1G0588300 | Bradi1g45020 | *HvPIN1a* (HORVU.MOREX.r3.7HG0666880) | O'Connor et al., Fusi et al.[46,47] |
| *BdPIN1b* | BdiBd21-3.3G0783600 | Bradi3g59520 | *HvPIN1/HvPIN1b* (HORVU.MOREX.r3.6HG0615550) | O'Connor et al., Kirschner et al., Fusi et al.[46–48] |
| *BdPME53-like* | BdiBd21-3.2G0255500 | Bradi2g19420 | *AtPME53* | Wu et al.[49] |
| *BdSCAP1* | BdiBd21-3.1G0206300 | Bradi1g15420 | *AtSCAP1* | Negi et al.[50] |
| *BdSCRM2* | BdiBd21-3.2G0762000 | Bradi2g59497 | *OsSCRM2* | Raissig et al., Wu et al.[21,30] |
| *BdSDD1-like* | BdiBd21-3.1G1009000 | Bradi1g75550 | *AtSDD1* | Von Groll et al.[51] |
| *BdSERK1/2-like1* | BdiBd21-3.5G0153700 | Bradi5g12227 | *AtSERK1/2* | Meng et al.[52] |
| *BdSERK1/2-like2* | BdiBd21-3.3G0620800 | Bradi3g46747 | *AtSERK1/2* | Meng et al.[52] |
| *BdSERK1/2-like3* | BdiBd21-3.3G0209300 | Bradi3g15660 | *AtSERK1/2* | Meng et al.[52] |
| *BdSLAH2* | BdiBd21-3.2G0203400 | Bradi2g15500 | *AtSLAH2* | Maierhofer et al.[53] |
| *BdSLP1-like1* | BdiBd21-3.3G0292100 | Bradi3g21000 | *SbSLP1* | Kumar et al.[54] |
| *BdSLP1-like2* | BdiBd21-3.3G0292000 | Bradi3g20980 | *SbSLP1* | Kumar et al.[54] |
| *BdSLP1-like3* | BdiBd21-3.3G0292700 | Bradi3g21030 | *SbSLP1* | Kumar et al.[54] |
| *BdSLP1-like4* | BdiBd21-3.5G0061500 | Bradi5g04630 | *SbSLP1* | Kumar et al.[54] |
| *BdSLP1-like5* | BdiBd21-3.1G0921800 | Bradi1g68400 | *SbSLP1* | Kumar et al.[54] |
| *BdSLP1-like6* | BdiBd21-3.1G0921700 | Bradi1g68390 | *SbSLP1* | Kumar et al.[54] |
| *BdSLP1-like7* | BdiBd21-3.3G0291900 | Bradi3g20970 | *SbSLP1* | Kumar et al.[54] |
| *BdSLP1-like8* | BdiBd21-3.3G0292200 | Bradi3g21010 | *SbSLP1* | Kumar et al.[54] |
| *BdSLP1-like9* | BdiBd21-3.1G0855400 | Bradi1g63370 | *SbSLP1* | Kumar et al.[54] |
| *BdSPCH1* | BdiBd21-3.1G0523400 | Bradi1g38650 | *AtSPCH, OsSPCH1* | Pillitteri and Torii, Raissig et al., Wu et al.[21,30,39] |
| *BdSPCH2* | BdiBd21-3.3G0131200 | Bradi3g09670 | *AtSPCH, OsSPCH2* | Pillitteri and Torii, Raissig et al., Wu et al.[20,21,30,39] |
| *BdSPL10* | BdiBd21-3.1G0421000 | Bradi1g31390 | *OsSPL10, ZmSPL14* | Lan et al., Kong et al.[55,56] |
| *BdSPL5* | BdiBd21-3.3G0074300 | Bradi3g05720 | *OsSPL5, ZmSPL10* | Xie et al., Kong et al.[56,57] |

| **Name** | **Bd21-3 accession** | **Bradi accession** | **Related gene in other species** | **Relevant literature** |
|---|---|---|---|---|
| *BdSTOMAGEN-1* | BdiBd21-3.2G0749400 | Bradi2g58540 | *AtSTOMAGEN, TaSTOMAGEN-1* | Sugano et al., Jangra et al.[12,58] |
| *BdTCP4-like* | BdiBd21-3.2G0089600 | Bradi2g06890 | *AtTCP4* | Palatnik et al.[59] |
| *BdTMM* | BdiBd21-3.2G0561000 | Bradi2g43940 | *AtTMM* | Geisler et al.[60] |
| *BdTMO6-like* | BdiBd21-3.1G0348000 | Bradi1g26570 | *AtTMO6* | Schlereth et al.[61] |
| *BdTSO1* | BdiBd21-3.1G0741600 | Bradi1g55710 | *AtTSO1* | Simmons et al.[62] |
| *BdVND-like1* | BdiBd21-3.1G1023300 | Bradi1g76732 | *AtVNDs* | Lehmann and Schneider[63] |
| *BdVND-like2* | BdiBd21-3.5G0221500 | Bradi5g16917 | *AtVNDs* | Lehmann and Schneider[63] |
| *BdWOX3B* | BdiBd21-3.2G0477300 | Bradi2g37650 | *OsWOX3B, ZmWOX3A* | Angeles-Shim et al., Li et al., Kong et al.[56,64–66] |
| *BdWOX4* | BdiBd21-3.5G0316500 | Bradi5g24080 | *OsWOX4* | Ohmori et al.[22] |
| *BdWOX9C-like1* | BdiBd21-3.2G0587500 | Bradi2g46055 | *AtWOX9* | Haecker et al.[67] |
| *BdXCP1-like* | BdiBd21-3.2G0501500 | Bradi2g39320 | *AtXCP1* | Funk et al.[68] |
| *BdYDA1* | BdiBd21-3.5G0238000 | Bradi5g18180 | *AtYDA, HvYDA1* | Bergmann et al., Abrash et al., Liu et al.[69–71] |
| *BdYDA2* | BdiBd21-3.3G0680900 | Bradi3g51380 | *AtYDA, HvYDA2* | Bergmann et al., Abrash et al., Liu et al.[69–71] |
| *DUF567* | BdiBd21-3.4G0612300 | Bradi4g44178 | N/A | This paper |
| *LRR kinase* | BdiBd21-3.2G0614100 | Bradi2g48000 | N/A | This paper |

**Table S4.** Seurat marker genes for all clusters

*see separate file "TableS4.xlsx"*

**Table S5.** Differentially expressed genes in non-protoplasted vs. protoplasted bulk RNA-seq

*see separate file "TableS5.xlsx"*

**Table S6.** Borda-ranked transcription factors per stomatal cluster (Mini-Ex analysis)

*see separate file "TableS6.xlsx"*

**Table S7.** Targetome of BdMUTE and BdFAMA in dividing GMC cluster

*see separate file "TableS7.xlsx"*
